# Seasonality and inter-annual stability in the population genetic structure of Batrachospermum gelatinosum (Rhodophyta)

**DOI:** 10.1101/2024.09.20.614195

**Authors:** Sarah Shainker-Connelly, Solenn Stoeckel, Morgan L. Vis, Roseanna M. Crowell, Stacy Krueger-Hadfield

## Abstract

Temporal population genetic studies have investigated evolutionary processes, but few have characterized the temporal patterns of reproductive system variation. Yet, temporal sampling may improve our understanding of reproductive system evolution through assessing the relative rates of selfing, outcrossing, and clonality. In this study, we focus on the monoicous, haploid-diploid freshwater red alga *Batrachospermum gelatinosum.* This species has a perennial, microscopic diploid phase (chantransia) that produces an ephemeral, macroscopic haploid phase (gametophyte). Recent work focusing on single time point genotyping suggested high rates of intragametophytic selfing, though there was variation among sites. We expand on this work by genotyping 191 gametophytes from four sites with reproductive system variation at multiple time points within and among years. Intra-annual data suggest shifts in gametophytic genotypes present throughout the gametophytic season. We hypothesize this pattern is likely due to the seasonality of the life cycle and the timing of meiosis among the chantransia. Inter-annual patterns were characterized by consistent genotypic and genetic composition, indicating stability in the prevailing reproductive system through time. Yet, our study identified limits to which available theoretical predictions and analytical tools can resolve reproductive system variation using haploid data. There is a need to develop better tools to understand the evolution of sex by expanding our ability to characterize the spatiotemporal variation in reproductive systems across diverse life cycles.

## INTRODUCTION

Traditionally, population genetic studies aim to characterize spatial genetic diversity and structure from samples collected at single time points from multiple sites (Storfer et al., 2007; Whitlock, 1992). Fewer studies have characterized temporal genetic diversity and structure by incorporating samples collected at multiple time points (Storfer et al., 2007; Whitlock, 1992). Yet, temporal genotyping has previously been used to address several types of ecological and evolutionary questions, including estimating population size (Waples, 1989), measuring evolutionary processes through time (Drummond et al., 2003), and assessing natural selection through time (e.g., from climate change, Jump et al., 2006; variable environmental conditions, Gómez et al., 1995; ecological succession, Linhart & Grant, 1996; or during biological invasions, Forsström et al., 2017). Though temporal variation likely also affects the prevailing reproductive mode, most studies assessing the reproductive system (i.e., the relative rates of sexual versus asexual and outcrossing versus selfing; Barrett, 2011) have only sampled a population once (e.g. Whitehead et al., 2018). Ecological variation, such as spatial and temporal fluctuation in pollinators (Barrett, 2015; Coates et al., 2013), can drive temporal variation in outcrossing rates within and among years (Whitehead et al., 2018), thereby influencing the partitioning of genetic diversity within and among populations. Additionally, in partially clonal organisms which can reproduce both sexually and asexually (often simultaneously), environmental conditions can result in different relative rates of sexual versus asexual reproduction through time (Bengtsson & Ceplitis, 2000; Gilabert et al., 2009; Liu et al., 2013; Weeks, 1993). For example, environmental stress can cause sexual reproduction in some aphid lineages (Simon et al., 2010), and seasonal changes can drive alternations of sexual and asexual reproduction in some microalgae (e.g., Dia et al., 2014; Lebret et al., 2012). Halkett et al. (2005) suggested using temporal sampling to monitor the evolution of clonal rates through time, but temporal studies are largely lacking. Thus, spatial and temporal sampling may be necessary to resolve the processes that drive reproductive mode variation (Becheler et al., 2017; Halkett et al., 2005) since the reproductive system shapes genetic diversity (Hamrick & Godt, 1996) and drives evolutionary responses to environmental change (e.g., Eckert et al., 2010; Orive et al., 2017).

Several reproductive system studies have characterized temporal variations in traditional population genetic summary statistics (e.g., genetic differentiation, genotypic diversity, inbreeding coefficients, and linkage disequilibrium) and how they vary with seasons and environmental conditions (Guillemaud et al., 2003, Tibayrenc & Ayala, 2012; Reynolds et al., 2017). For example, some aphids are cyclical parthenogens in which there are periods of asexual reproduction followed by a sexual event, resulting in some genetic variability within years but stability between years (Guillemaud et al., 2003, 2011). This phenomenon is distinct from other types of partial clonality in which sexual and asexual reproduction occur simultaneously. For example, genetic differentiation between years highlighted the importance of the banking and continuous germination of resting cysts on the bloom dynamics of microalgae (Dia et al., 2014; Lebret et al., 2012). For diploids, linkage disequilibrium, heterozygosity, and the inbreeding coefficient *F_IS_* have often been used as proxies to estimate reproductive modes and their temporal variations (Arnaud-Haond et al., 2007; Allen & Lynch, 2012; Stoeckel & Masson, 2014; Bürkli et al., 2017), though they can be inaccurate for low to moderate clonal rates and cannot be assessed in the haploid phase. While these comparative methods can be informative, traditional population genetic summary statistics do not provide direct rates of clonality and selfing.

Reproductive modes can be more directly characterized using dedicated methods to estimate changes in genotypic frequencies through time (Becheler et al., 2017; Stoeckel et al., 2024).

These methods require three conditions that can be difficult to meet for non-model species with complex life cycles: (i) prior knowledge of the generation time of natural populations, as samples should be collected one generation apart, (ii) prior knowledge of inbreeding and selfing rates, and (iii) diploid genotypes. Therefore, these methods are difficult, or in some cases impossible, to use for haploid-diploid organisms, such as algae, ferns, and some fungi (Becheler et al., 2017). Yet, macroalgae present an opportunity to better understand the evolution of sex because many exhibit spatial and temporal separation of meiosis and fertilization resulting in two phases – often haploid and diploid – of long duration (Otto & Marks, 1996). Algae exhibit a broad diversity of life cycle types, are phylogenetically diverse, and have thus been proposed for testing the influence of life cycles on reproductive system variation (see Krueger-Hadfield et al., 2021; Krueger-Hadfield, 2024; Olsen et al., 2020; Otto & Marks, 1996). Most previous data have been taxonomically restricted to brown algae (see Heesch et al., 2021) and marine red algae (e.g., Engel et al., 1999, 2004; Krueger-Hadfield et al., 2013, 2015; Guillemin et al., 2008; see additionally Krueger-Hadfield et al., 2021). Otto & Marks (1996) proposed green algae as useful models for testing predictions for reproductive system variation across life cycle types, but there are still too few data to accurately test these hypotheses (see Krueger-Hadfield 2024).

We recently expanded the taxonomic breadth to include freshwater red algae (Krueger-Hadfield et al. 2024). Not only do they display variation in their sexual systems (i.e., monoicy and dioicy; see Figure 2 in Krueger-Hadfield et al. 2024), but species in the Batrachospermales have a unique haploid-diploid life cycle in which a microscopic, perennial diploid phase – called the chantransia – alternates with a macroscopic, ephemeral haploid gametophyte phase. Unlike marine red algae, the gametophytes remain physically attached to the chantransia. This life cycle results in unique eco-evolutionary consequences for these types of red algae (see Krueger-Hadfield et al., 2024; see Figure 1 in Shainker-Connelly et al., 2024). First, intragametophytic selfing describes fertilization between a spermatium (sperm) and a carpogonium (egg) produced by the same monoicous gametophyte, and results in instantaneous, genome-wide homozygosity, unlike selfing in diploid-dominant or dioicous organisms (Klekowski, 1969; see Figure 1a in Shainker-Connelly et al., 2024). This type of selfing is uncommon in marine red algae, most of which are dioicous (but see the following exceptions – Fujio et al., 1985; Maggs, 1988; Lindstrom, 1993). Second, intergametophytic selfing describes fertilization that occurs between two gametophytes originating from the same parental chantransia and is expected to gradually erode genetic diversity in a population, similarly to selfing in diploid-dominant organisms (Klekowski, 1969a). Unlike the gametophytes and tetrasporophytes of marine red algae, the batrachospermalean gametophyte remains physically attached to the chantransia, and it is possible that different filaments of the same chantransia could produce multiple gametophytes. The proximity of the resulting gametophytes could facilitate intergametophytic selfing. Though arising from the physical attachment of multiple gametophytes to the same chantransia, the resulting genetic pattern may be analogous to the clumped dispersal of tetraspores in *Chondrus crispus* (see Krueger-Hadfield et al. 2013, 2015) which has been proposed to drive a pattern of discrete genotypes in close proximity to one another (Krueger-Hadfield, 2011). Finally, the chantransia can reproduce asexually by producing monospores, which germinate and develop into new chantransia (Sheath, 1984; see Figure 1d in Shainker-Connelly et al., 2024). The frequency of monospore production in natural populations is unknown. Moreover, it would be difficult to distinguish between monospore production by a chantransia and different carpospores from the same gametophyte germinating into chantransia as both would result in many chantransia sharing the same genotype. The life cycle highlights the complications that arise when characterizing the reproductive systems of non-model, haploid-diploid taxa, and the necessity of expanding our toolbox to encompass eukaryotic diversity more broadly.

**Figure 1.**
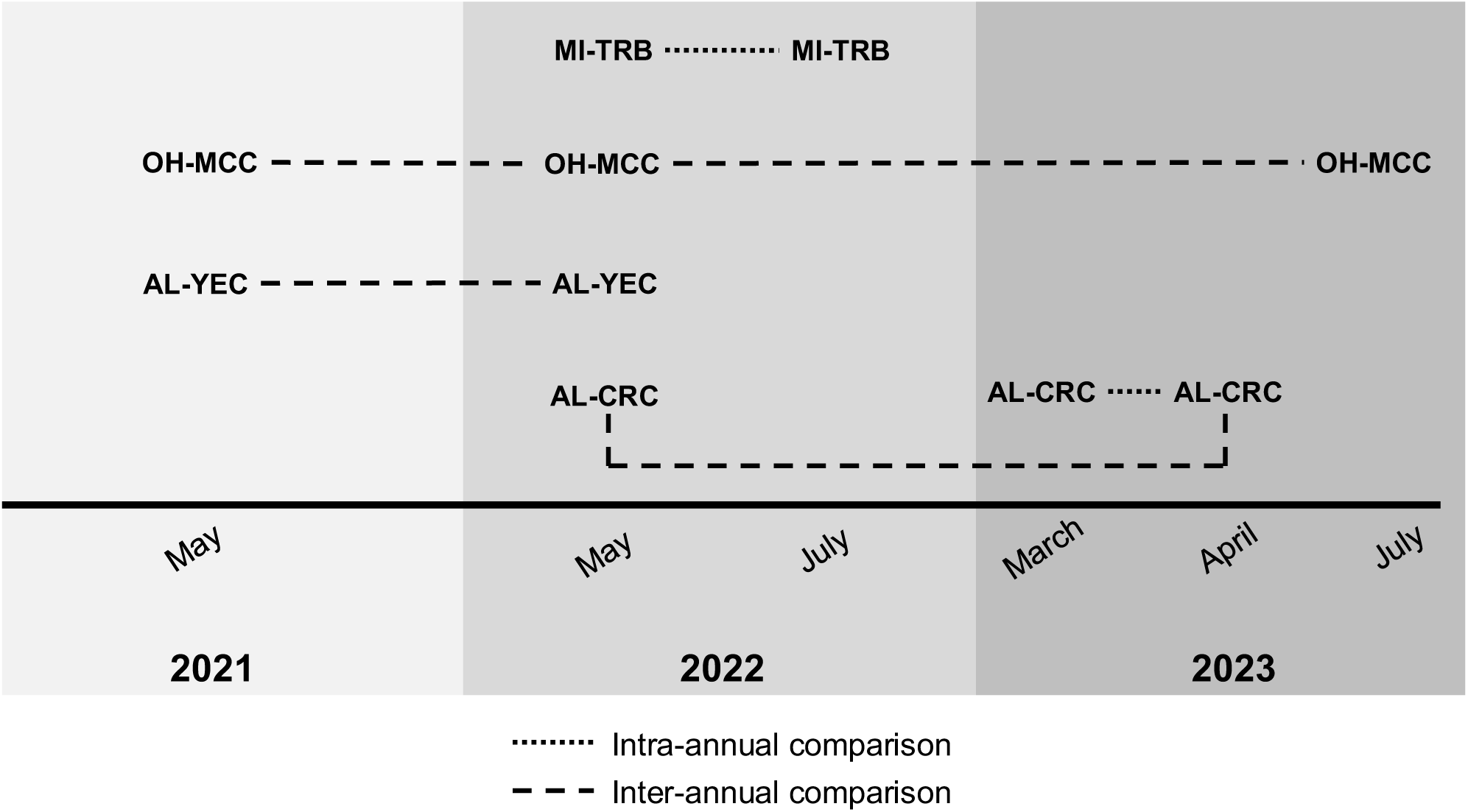
Conceptual diagram for the sampling dates at each site, shown in latitudinal order from north to south. Months and years are indicated along the horizontal axis. Shading distinguishes each year. Traverse River on Big Traverse Rd., MI (MI-TRB) was sampled in May 2022 and July 2022 for an intra-annual comparison. Monday Creek, OH (OH-MCC) was sampled in May 2021, May 2022, and July 2023, providing inter-annual comparisons across three years. Yellow Creek, AL (AL-YEC) was sampled in May 2021 and May 2022, for an inter-annual comparison between two years. Cripple Creek, AL (AL-CRC) was sampled in May 2022, March 2023, and April 2023, for both an inter-annual comparison and an intra-annual comparison.

**Figure 2.**
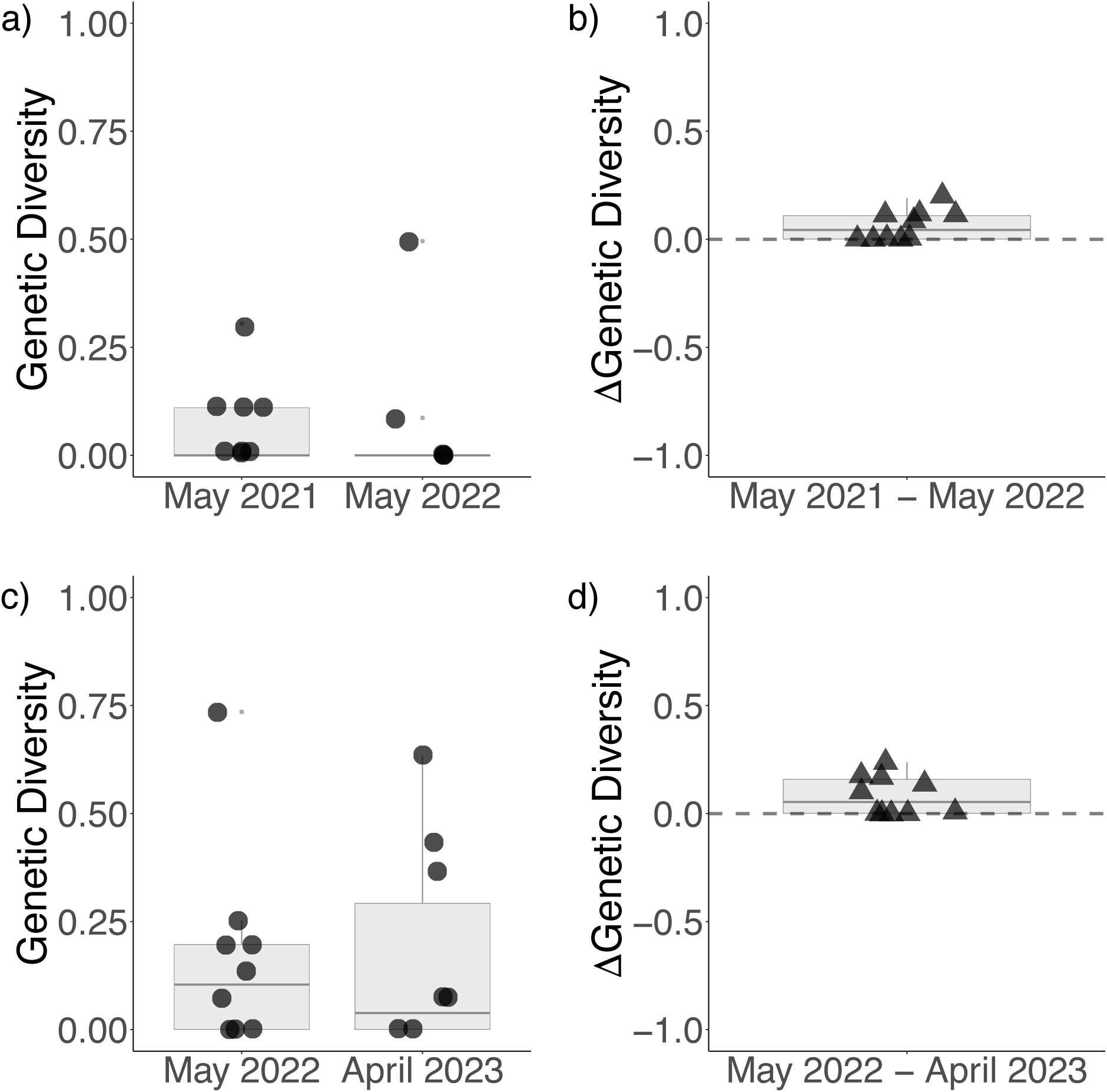
The distribution of genetic diversity (calculated as expected heterozygosity, *H_E_*) for each locus for inter-annual time points at a) Yellow Creek (AL-YEC) and c) Cripple Creek (AL- CRC). The y-axis range is shown as 0 to 1 for genetic diversity values in a) and c). Changes in genetic diversity (shown as ΔGenetic Diversity) between years for each locus are shown for b) Yellow Creek (AL-YEC) and d) Cripple Creek (AL-CRC). The y-axis range is shown as -1.0 to 1.0 for the change in genetic diversity estimates per locus in b) and d), with a dashed grey line to indicate the y-intercept at 0. Boxes represent the interquartile range, the middle lines are medians, whiskers represent the 1.5 interquartile ranges, and the small light grey dots represent outliers.

We recently characterized the reproductive system of the monoicous, widespread red alga *Batrachospermum gelatinosum* (Sheath & Cole, 1992; Vis & Necchi Jr, 2021) across eastern North America (Shainker-Connelly et al., 2024). We interpreted the population genetic patterns we observed as indicative of high rates of intragametophytic selfing, though we noted that we could not distinguish between the genetic effects of intragametophytic selfing and monospore production using haploid gametophytic genotypes. Here, we expand on our previous work by using temporal genotyping to improve our understanding of the reproductive system of *B. gelatinosum.* Four of the sites from our previous study (Shainker-Connelly et al., 2024) were sampled at multiple time points to assess temporal patterns of reproductive system variability (Figure 1, Figure S1). From previous single snapshot genotyping, we observed genetic signatures of high rates of intragametophytic selfing at two sites, whereas the other two sites had more intermediate selfing rates (Shainker-Connelly et al., 2024).

We aimed to use inter-annual population genetic comparisons to better understand the temporal dynamics of the reproductive system (Figure 1). If a site has low levels of standing diversity maintained by high rates of intragametophytic selfing, we predicted that over time, (i) high linkage disequilibrium, (ii) low genetic diversity, (iii) low genotypic richness and evenness, (iv) low numbers of raw alleles per locus, (v) low number of diverging alleles between gametophyte pairs at all time points, and (vi) if genotypes remain the same, then there will be low genetic differentiation or if different genotypes become dominant in different years, then we expect high differentiation. If a population has intermediate to high levels of standing genetic diversity and begins to undergo greater rates of intragametophytic selfing, we predicted the following over time: (i) linkage disequilibrium will increase, (ii) genetic diversity will decrease, (iii) the raw number of alleles per locus and number of diverging alleles between pairs of gametophytes will decrease, and (iv) intermediate to high temporal genetic differentiation. If a population has intermediate levels of intragametophytic selfing mixed with intergametophytic selfing and outcrossing (i.e., mixed mating, Goodwillie et al., 2005), we predicted (i) high variance in single locus values of linkage disequilibrium, genetic diversity, and temporal genetic differentiation, and (ii) a broad distribution in the number of raw alleles per locus and the number of diverging alleles between gametophytic pairs (Table 1).

**Table 1.** Summary of temporal comparisons and predictions of potential outcomes for *Batrachospermum gelatinosum*.

| Comparison | Purpose | Sites and time points | Predictions | Genetic patterns |
| --- | --- | --- | --- | --- |
| Inter-annual | Temporal patterns in the reproductive system | <u>OH-MCC</u><br>May 2021<br>May 2022<br>July 2023<br><u>AL-YEC</u><br>May 2021<br>May 2022<br><u>AL-CRC</u><br>May 2022<br>April 2023 | High levels of intragametophytic selfing | *LOW DIVERSITY*<br>• *Linkage disequilibrium*: High with low variance<br>• *Genetic diversity*: Consistently low<br>• *Temporal genetic differentiation*: Low<br>• *Genotypic richness and evenness*: Low<br>• *Raw and diverging alleles*: Low |
|  |  |  | Increasing intragametophytic selfing from one year to the next | *INTERMEDIATE DIVERSITY*<br>• *Linkage disequilibrium*: Increase from year to year<br>• *Genetic diversity*: Decrease from year to year<br>• *Temporal genetic differentiation*: Intermediate to high<br>• *Raw and diverging alleles*: Decrease from year to year |
|  |  |  | Mixed mating system* | *HIGH DIVERSITY*<br>• *Linkage disequilibrium*: Low with high variance<br>• *Genetic diversity*: Intermediate to high with high variance<br>• *Temporal genetic differentiation*: Intermediate to high with high variance<br>• *Raw and diverging alleles*: Broad distribution |
| Intra-annual | Seasonality of the life cycle | <u>MI-TRB</u><br>May 2022<br>July 2022<br><u>AL-CRC</u><br>March 2023<br>April 2023 | All gametophytes at the same time of year | *CONSISTENT DIVERSITY*<br>• *Genetic diversity*: Consistent<br>• *Temporal genetic differentiation*: Low<br>• *Genotypic richness and evenness*: Consistent<br>• *Genotypes*: Similar between time points |
|  |  |  | Different chantransia genotypes undergo meiosis at different times to produce gametophytes throughout the season | *INCREASING DIVERSITY*<br>• *Genetic diversity*: Increase from one time point to the next<br>• *Temporal genetic differentiation*: High<br>• *Genotypic richness and evenness*: Increase from one time point to the next<br>• *Genotypes*: Transition between time points |

We aimed to use intra-annual comparisons to better understand the seasonality of the haploid-diploid *Batrachospermum gelatinosum* life cycle (Figure 1). The perennial chantransia likely enhances population stability by persisting through disturbances and maintaining abundance in the upstream reaches of drainage basins (Hambrook & Sheath, 1991). The macroscopic gametophytes typically appear seasonally, although they may be perennial in some cases (Sheath & Vis, 2015). If different chantransia (i.e., diploid individuals) produce gametophytes at the same time, then we predicted the following over time: (i) genetic diversity will remain consistent regardless of the time of year and (ii) there will be little if any genetic differentiation between intra-annual time points. If different chantransia produce gametophytes at different times during the season, we predicted the following over time: (i) shifts in genotypic composition, (ii) intermediate to high temporal genetic differentiation, and (iii) if more chantransia produce gametophytes later compared to earlier in the season, an increase in genetic diversity (Table 1). Our work provides a reference point for understanding the life cycle dynamics and temporal patterns of reproductive system variation in *Batrachospermum gelatinosum*, thereby expanding our understanding of eukaryotic reproductive system variation.

Moreover, this study is one of a handful of attempts to use temporal sampling to characterize population genetic parameters in natural populations.

## MATERIALS & METHODS

### Sample Collection

We collected *Batrachospermum gelatinosum* gametophytic thalli (hereafter referred to as gametophytes) from four sites in the eastern United States in 2021, 2022, and 2023 (Table 1, Figure S1). We did not sample the chantransia because they are microscopic and genotyping would be a challenge (see Schoenrock et al., 2020 for a discussion on microscopic forms). Sites AL-CRC (Alabama) and MI-TRB (Michigan) were each sampled twice within the same season to provide an intra-annual comparison of genotypes: AL-CRC in March 2023 and April 2023 and MI-TRB in May 2022 and July 2022. Sites AL-CRC, AL-YEC (Alabama), and OH-MCC (Ohio) were each sampled during sequential years, providing inter-annual comparisons of genotypes: AL-CRC in May 2022 and April 2023, AL-YEC May 2021 and May 2022, and OH-MCC in May 2021, May 2022, and July 2023 (Table 2, Figure 1). The sampling protocols were also described in Shainker-Connelly et al. (2024) and Crowell et al. (2024a). For each site, we used Google Maps or the iPhone app GPSCoordinates ver. 5.18 (Neal, 2018) to note GPS coordinates. We used a transect tape to measure stream width and length of the sampled reach, chosen based on accessibility and the presence of gametophytes. We visually estimated stream bed composition, water color, and water clarity near the middle of the sampling area (Table S1). At sites AL-CRC, AL-YEC, and MI-TRB (see Table 1), we used an Oakton PCTSTestr 50 Pocket Tester to measure pH, water temperature, and specific conductivity. At OH-MCC, we used an Oakton pHTestr 5 to measure pH and water temperature and an Oakton ECTestr low to measure specific conductivity. We used a flow probe (Global Water Instruments, Model FP111) to measure current velocity and stream depth at sites AL-CRC, AL-YEC, and MI-TRB (see Table 1). At OH-MCC, we used a different flow probe (General Oceanics, Mechanical Flow Meter) and measured stream depth with a ruler. At sites AL-CRC, AL-YEC, and MI-TRB (see Table 1), we used a spherical densiometer (Forest Densiometer, Model A) to calculate percent canopy cover following Lemmon (1956) and Lemmon (1957), but with one reading instead of four. At OH-MCC, canopy cover was estimated by eye.

**Table 2.** Locations and physiochemical data measured at each site in which *Batrachospermum gelatinosum* gametophytes were sampled at multiple time points. Dashes (-) indicate that the measurement was not taken.

| Site Name | Site | State | Sampling | Latitude | Longitude | Herbarium | pH | Specific | Water | Current | Stream | Stream | Sampling | Canopy |
| --- | --- | --- | --- | --- | --- | --- | --- | --- | --- | --- | --- | --- | --- | --- |
|  | Abbreviation |  | Date |  |  | Voucher |  | Conductivity | Temperature | Velocity | Depth | Width (m) | Length (m) | Cover (%) |
|  |  |  |  |  |  |  |  | (μS · cm-1) | (°C) | (m · s<sup>-1</sup>) | (cm) |  |  |  |
| Traverse River on | MI-TRB | Michigan | 12-May-2022 | 47.195156 | -88.239225 | BHO: A-1889 | 7.14 | 30.9 | 9.1 | 0.5 | 54 | 7.74 | 9.44 | 5.3 |
| Big Traverse Rd. |  |  | 26-Jul-2022 |  |  | BHO: A-1887 | - | - | - | - | - | - | - | - |
| Monday Creek | OH-MCC | Ohio | 19-May-2021 | 39.500500 | -82.246300 | BHO: A-1723 | 6.62 | 500 | 17.1 | - | 42 | 11.1 | 18 | 50 |
|  |  |  | 5-May-2022 |  |  | BHO: A-1814 | 8.0 | 510 | 16.2 | 0.3 | 70 | 7 | 40 | 20 |
|  |  |  | 23-Jul-2023 |  |  | BHO: A-1993 | 7.3 | 580 | 21.2 | 0.18 | 38 | 8.3 | 18 | 90 |
| Yellow Creek | AL-YEC | Alabama | 19-May-2021 | 33.572000 | -87.403000 | BHO: A-1902 | 7.06 | 151 | 19.5 | 0.4 | 50 | 5.65 | 33.6 | 18.7 |
|  |  |  | 2-May-2022 |  |  | BHO: A-2038 | 7.15 | 50 | 18.5 | 0.8 | 26 | 5.01 | 34.8 | - |
| Cripple Creek | AL-CRC | Alabama | 2-May-2022 | 33.492530 | -87.562633 | BHO: A-2037 | 7.34 | 124 | 16.5 | 0.3 | 17 | 6.5 | 45.16 | 64 |
|  |  |  | 16-Mar-2023 |  |  | BHO: A-2064 | 7.41 | 86.3 | 7.5 | 0.6 | 31 | 7.3 | 48 | 13 |
|  |  |  | 20-Apr-2023 |  |  | BHO: A-2065 | 7.22 | 129.6 | 13 | 0.6 | 17 | 7.5 | 48 | 39 |

We haphazardly sampled gametophytes within the measured sampling length of the reach (see Table 2), aiming to collect 20-30 gametophytes from each time point at each site. At some time points, there were few gametophytes with a patchy distribution, resulting in a smaller sample size (Table 3). The stream we sampled at AL-CRC contains three distinct sections: an upstream riffle, a pool, and a downstream riffle. We took environmental measurements within each of these three distinct sections and noted the sections from which gametophytes were collected. Most *Batrachospermum gelatinosum* gametophytes were collected in the riffles, so the environmental measurements taken in the upstream riffle section are reported, except for the sampling length which includes both riffles and the pool (Table 2).

**Table 3.** Summary statistics calculated for sites in which *Batrachospermum gelatinosum* gametophytes were sampled at multiple time points, with site abbreviations indicated (see Table 1). The third column (“Intra” or “inter”) indicates whether the time point was used for an inter-annual comparison, an intra-annual comparison, or both. Ten microsatellite loci were used for all sites except OH-MCC, for which nine microsatellite loci were used because locus Bgel_071 did not amplify. Summary statistics include the following: n, total number of gametophytes genotyped; *G*, number of unique genotypes; *pid*, probability of identity of sibs; *R*, genotypic richness; *D\**, genotypic evenness; *pareto ß*, distribution of clonal membership, *H*_E_, expected heterozygosity *H_E_*, multilocus estimate of linkage disequilibrium; and the number of fixed loci per sampling date.

| Site | Sampling | n | G | *pid* | *R* | *D\** | *H<sub>E</sub>* | Pareto *β* | $\bar{r}_d$ | Number of | Mean allelic |
| --- | --- | --- | --- | --- | --- | --- | --- | --- | --- | --- | --- |
| Abbreviation | Date |  |  |  |  |  |  |  |  | fixed loci | richness |
| *Inter-annual comparisons* |  |  |  |  |  |  |  |  |  |  |  |
| OH-MCC | 19-May-2021 | 20 | 1 | 1 | 0 | 0 | 0 | 0.02 | 1 | 9 | 1.00 |
|  | 5-May-2022 | 28 | 2 | 0.666 | 0.04 | 0.07 | 0.015 | 0.01 | 1 | 7 | 1.15 |
|  | 23-Jul-2023 | 19 | 1 | 1 | 0 | 0 | 0 | 0.02 | 1 | 9 | 1.00 |
| AL-YEC | 19-May-2021 | 17 | 5 | 0.150 | 0.25 | 0.43 | 0.064 | 0.10 | 0.29 | 7 | 1.20 |
|  | 2-May-2022 | 22 | 3 | 0.285 | 0.10 | 0.56 | 0.058 | 0.16 | 0.03 | 8 | 1.13 |
| AL-CRC | 2-May-2022 | 27 | 15 | 0.004 | 0.54 | 0.88 | 0.159 | 0.49 | 0.04 | 4 | 1.49 |
|  | 20-Apr-2023<sup>a</sup> | 25 | 11 | 0.012 | 0.42 | 0.89 | 0.159 | 0.66 | 0.04 | 5 | 1.42 |
| *Intra-annual comparisons* |  |  |  |  |  |  |  |  |  |  |  |
| MI-TRB | 12-May-2022 | 6 | 4 | 2.17E-05 | 0.60 | 0.80 | 0.344 | 0.63 | 0.66 | 2 | 2.30 |
|  | 26-Jul-2022 | 13 | 6 | 4.76E-05 | 0.42 | 0.64 | 0.308 | 0.23 | 0.50 | 2 | 1.97 |
| AL-CRC | 16-Mar-2023 | 14 | 8 | 0.002 | 0.54 | 0.89 | 0.190 | 0.84 | 0.20 | 4 | 1.60 |
|  | 20-Apr-2023<sup>a</sup> | 25 | 11 | 0.012 | 0.42 | 0.89 | 0.159 | 0.66 | 0.04 | 5 | 1.42 |

<sup>a</sup>Samples from this time point were used for both inter- and intra-annual comparisons.

**Table 4.** The raw number of alleles counted at each locus, for each sampling point, is provided. A dash (-) is entered for locus Bgel_071 at all time points at OH-MCC since this locus did not amplify for the gametophytes sampled at this site.

| Site | Sampling |  |  |  |  |  |  |  |  |  |  |
| --- | --- | --- | --- | --- | --- | --- | --- | --- | --- | --- | --- |
| Abbreviation | Date | Bgel_021 | Bgel_071 | Bgel_052 | Bgel067 | Bgel_053 | Bgel_070 | Bgel_059 | Bgel_073 | Bgel_057 | Bgel_056 |
| MI-TRB | 12-May-2022 | 1 | 4 | 2 | 1 | 3 | 2 | 2 | 2 | 3 | 3 |
|  | 26-Jul-2022 | 1 | 4 | 2 | 1 | 3 | 5 | 2 | 2 | 3 | 3 |
| OH-MCC | 19-May-2021 | 1 | - | 1 | 1 | 1 | 1 | 1 | 1 | 1 | 1 |
|  | 5-May-2022 | 2 | - | 2 | 1 | 1 | 1 | 1 | 1 | 1 | 1 |
|  | 23-Jul-2023 | 1 | - | 1 | 1 | 1 | 1 | 1 | 1 | 1 | 1 |
| AL-YEC | 19-May-2021 | 2 | 3 | 2 | 1 | 1 | 2 | 1 | 1 | 1 | 1 |
|  | 2-May-2022 | 1 | 2 | 1 | 1 | 1 | 1 | 1 | 1 | 1 | 2 |
| AL-CRC | 2-May-2022 | 1 | 6 | 1 | 2 | 2 | 2 | 1 | 1 | 2 | 2 |
|  | 16-Mar-2023 | 1 | 4 | 1 | 2 | 3 | 2 | 1 | 1 | 2 | 2 |
|  | 20-Apr-2023 | 1 | 6 | 1 | 2 | 2 | 2 | 1 | 1 | 1 | 2 |

We used a dissecting microscope (40X magnification) to look for carposporophytes on each gametophyte. In our previous work (Crowell et al., 2024a; Shainker-Connelly et al., 2024; Crowell et al., 2024b), the presence of carposporophytes did not result in ‘diploid’ gametophytes with two or more alleles. This could be due to high rates of intragametophytic selfing in which the same gametophyte produced both the carpogonium (i.e., egg) and spermatium (i.e., sperm) that resulted in the formation of the carposporophyte. Alternatively, DNA from the genotyped gametophyte may have swamped any exogenous paternal DNA in the carposporophytes. We ensured that we preserved a single gametophyte by physically separating thalli if entangled with one another and by removing the lower portion of the thallus to ensure that there were no remnants of the chantransia. We used silica gel (Activa Flower Drying Art Silica Gel) to preserve tissue from each gametophyte. When possible, the remaining thallus was mounted on herbarium paper (University of California-type Herbarium Mounting Paper, Herbarium Supply, Bozeman, MT), and representative vouchers were deposited in the Floyd Bartley Herbarium at Ohio University (Table 2).

### DNA Extraction and PCR Amplification

We extracted total genomic DNA using the Machery-Nagel Nucleospin® Plant II kit (Macherey-Nagel) following the manufacturer’s protocol, except that the lysate was incubated at room temperature for one hour and DNA was eluted in a 100 μL volume of molecular grade water (see also Crowell et al., 2024a; Shainker-Connelly et al., 2024).

We used ten previously developed microsatellite loci (locus development described in Crowell et al., 2024b; phylogeographic patterns described in Crowell et al., 2024a; reproductive system variation from one shot genotyping described in Shainker-Connelly et al., 2024). We amplified most loci for most gametophytes using multiplex PCRs with a final volume of 15 μL: 2 μL of DNA, forward and reverse primers (Table S2), 1X Promega GoTaq® Flexi Buffer (Promega, Madison, WI, USA; Cat #M890A), 2 mM of MgCl_2_ (Promega, Madison, WI, USA; Cat #A351H), 250 μM of each dNTP (Promega, Madison, WI, USA; Cat #R0192), 1mg/mL of BSA, and 1.0 U of Promega GoTaq® Flexi DNA Polymerase (Promega, Madison, WI, USA; Cat #M829B). For the locus Bgel_056, we used simplex PCRs with a final volume of 15 μL: 2 μL of DNA, 150 nM of the forward labeled primer, 100 nM of the forward unlabeled primer, 250 nM of the unlabeled reverse primer, 1X buffer (Promega, Madison, WI, USA; Cat #M890A), 2 mM of MgCl_2_, 250 μM of each dNTP (Promega, Madison, WI; Cat #R0192), 1mg/mL of BSA, and 1U of Promega GoTaq® Flexi DNA Polymerase (Promega, Madison, WI, USA; Cat #M829B). For reruns of any locus that did not amplify in the first attempt in a multiplex PCR, we used simplex PCRs with a final volume of 15 μL: 2 μL of DNA, 250 nM of the forward labeled primer, 250 nM of the unlabeled reverse primer, 1X buffer (Promega, Madison, WI, USA; Cat #M890A), 2 mM of MgCl_2_, 250 μM of each dNTP (Promega, Madison, WI; Cat #R0192), 1mg/mL of BSA, and 1U of Promega GoTaq® Flexi DNA Polymerase (Promega, Madison, WI, USA; Cat #M829B). We used the following PCR program: 95°C for 2 minutes, followed by 35 cycles of 95°C for 30 seconds, 59°C for 30 seconds, and 72°C for 30 seconds, with a final extension stage of 72°C for 5 minutes.

We diluted 1.5 μL PCR product in 9.7 μL HiDi formamide (Applied Biosystems, Cat #4311320) and 0.30 μL GS 500 LIZ (Applied Biosystems, Cat #4322682). Fragment analysis was then performed at the Heflin Center for Genomic Sciences at the University of Alabama at Birmingham. We scored alleles using Geneious Prime v.2022.2.2 (https://www.geneious.com) then manually checked each bin with those previously described in Crowell et al. (2024b), adjusting when the raw allele size was slightly larger or smaller than the previously defined bins (Table S3).

After several PCR attempts, locus Bgel_071 did not amplify for most gametophytes sampled for all three time points at OH-MCC. This result was consistent with a geographic pattern previously detected at this site (Crowell et al., 2024a; Shainker-Connelly et al, 2024). Bgel_071 was removed from OH-MCC gametophytes so that a total of nine loci were used for analyses of all time points at this site.

### Data Analyses

Gametophytes for which any loci did not amplify after several PCR attempts were excluded from subsequent analyses. The null allele frequency was directly estimated by calculating the percent of gametophytes that did not amplify at each locus after several PCR attempts and after discounting technical errors (see also Krueger-Hadfield et al., 2011).

We calculated the following multilocus summary statistics for each time point at each site to describe the reproductive system, following the recommendations and methods described by Stoeckel et al. (2021) and implemented in Krueger-Hadfield et al. (2021). We calculated the probability of identity between sibs (*pid*), which ranges from 0-1, to assess whether loci were of sufficient resolution to distinguish individuals (Jacquard, 2012; Waits et al., 2001). Then, we calculated genotypic richness (*R*), which provides information on the relative proportion of repeated multilocus genotypes (MLGs), as: 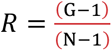 where G is the number of distinct genotypes (i.e., genets) and *N* is the number of genotyped gametophytes (Dorken & Eckert, 2001). We also calculated genotypic evenness, which provides information about the relative abundance of each MLG in a site (*D\**, see Box 3 in Arnaud-Haond et al., 2007), and is expected to increase with increasing *R* in outcrossed populations (Baums et al., 2006; Krueger-Hadfield et al., 2021).

We calculated multilocus and per-locus values of expected heterozygosity (*H*_E_) following Stoeckel et al. (2021). To calculate mean allelic richness for each time point, we used the allelic.richness function in the R package hierfstat (Goudet & Jombart, 2015). We then described the distribution of clonal membership (*pareto ß*) for each time point sampled (Table 3). As *Batrachospermum gelatinosum* is monoicous, intragametophytic selfing results in similar population genetic patterns as asexual reproduction via monospores by the chantransia, so pareto *β* cannot be used to disentangle the effects of these two reproductive modes (see Shainker- Connelly et al., 2024). We considered that pareto *β* > 2 was associated with low rates of intragametophytic selfing, 0.7 < pareto *β* < 2 was associated with intermediate rates of intragametophytic selfing, and pareto *β* < 0.7 was associated with high rates of intragametophytic selfing following similar predictions for asexual reproduction based on empirical data in Krueger-Hadfield et al. (2021).

For each time point at each site, we calculated multilocus values of linkage disequilibrium (*r̄_d_*) following Agapow & Burt (2001). We also determined linkage disequilibrium (|D’|) between each pair of alleles for each time point (Lewontin, 1964). In partially clonal taxa, including red macroalgae, both asexual reproduction and selfing can lead to an increase in *r̄_d_* and pairwise |D’| values, and variance of these values, within a species (Krueger-Hadfield et al., 2021; Stoeckel et al., 2021).

To measure genetic differentiation between time points at the same site, we calculated a pairwise measure of temporal genetic differentiation, like *F_ST_*, for each locus. This value ranges from 0 - 1, where 0 refers to no differentiation and 1 refers to fixation of different alleles.

Finally, we calculated raw genetic distances for all pairs of gametophytes using GenAPoPop (Stoeckel et al., 2024) adapted for haploid data. The maximum possible number of diverging alleles between a pair of gametophytes was ten for all sites except OH-MCC. Since Bgel_071 was excluded for OH-MCC, the maximum possible number of diverging alleles between pairs of gametophytes at all time points was nine at OH-MCC. We calculated the number of diverging alleles between all pairs of gametophytes for each site within each sampled time point. Finally, we calculated the number of raw alleles present at each locus.

### Data Visualization

Figures were prepared using R ver. 2022.07.2 (R Core Team, 2022) with the following packages: ggplot2 (Wickham, 2016), gridExtra (Auguie, 2017), pastecs (Grosjean & Ibanez, 2018), and car (Fox & Weisberg, 2019).

## RESULTS

### Genotyping and null alleles

We generated a total of 191 gametophytic genotypes. We attempted to amplify all loci for 208 gametophytes from four sites collected at ten different time points. One gametophyte did not amplify at locus Bgel_021, five at Bgel_067, one at Bgel_053, four at Bgel_059, and ten at Bgel_057. All gametophytes amplified at loci Bgel_052, Bgel_070, Bgel_073, and Bgel_056.

Locus Bgel_071 displayed a geographic pattern of non-amplification, in which 65 out of 68 gametophytes from OH-MCC (Ohio) had what we presume are one or more null alleles, possibly due to an insertion or deletion in the sequence between the locus-specific primers (Crowell et al., 2024a; Crowell et al., 2024b; Shainker-Connelly et al., 2024). At locus Bgel_071, null allele frequency reached 31.3% when OH-MCC was included, but all gametophytes from other sites amplified at this locus (Table S4). Excluding Bgel_071, null allele frequencies were low overall (less than 4.8%; Table S4). After 17 gametophytes with one or more loci that did not amplify were removed, 191 gametophytic genotypes remained. The probability of identity (*pid*) was -7.55 x 10^-9^ over all 124 samples genotyped with ten loci (sites AL-CRC, AL-YEC, Alabama; and MI-TRB, Michigan), and 0.842 over all 67 samples genotyped with nine loci (site OH-MCC).

### Inter-annual comparisons

Among time points, *pid* values remained stable at a site. At OH-MCC, *pid* ranged from 0.666 in May 2022 to 1 in May 2021 and July 2023. At AL-YEC, *pid* ranged from 0.150 in May 2021 to 0.285 in May 2022. At AL-CRC, *pid* ranged from 0.004 in May 2022 to 0.012 in April 2023 (Table 3).

Genotypic richness and evenness varied among sites but remained stable between inter- annual time points sampled from a site. At OH-MCC, genotypic richness (*R*) ranged from 0.00 to 0.04, while genotypic evenness (*D\**) ranged from 0.00 to 0.07 (Table 3). One multilocus genotype (MLG) was shared among all time points and one MLG was represented by a single gametophyte in May 2022. At AL-YEC, *R* decreased slightly from 0.25 in May 2021 to 0.10 in May 2022, while *D\** increased slightly from 0.43 in May 2021 to 0.56 in May 2022 (Table 3).

One MLG was shared among 13 and 9 gametophytes from May 2021 and May 2022, respectively. Four unique MLGs were represented in May 2021, and one unique MLG in May 2022. At AL-CRC, *R* decreased slightly from 0.54 in May 2022 to 0.42 in April 2023, while *D\** remained stable at 0.88 in May 2022 and 0.89 in April 2023 (Table 3). Six MLGs were repeated between years, nine MLGs were unique to May 2022, and four MLGs to April 2023.

Among inter-annual time points at the same site, genetic diversity, measured as expected heterozygosity (*H_E_*), was stable and low (Figure 2). At OH-MCC, *H_E_* ranged from 0.000 to 0.015. Single locus *H_E_* at this site were all 0.000 in May 2021 and July 2023. In May 2022, all loci except two (with *H_E_* = 0.069) had genetic diversity values of 0.000, for a mean and standard error of 0.015 ± 0.010 at that time point (Table 3). Due to these two loci, the change in *H_E_* ranged from -0.069 to 0.069 over both intervals, while the remaining seven loci remained consistent. At AL-YEC, *H_E_* was 0.064 in May 2021 and 0.058 in May 2022. The single locus genetic values ranged from 0.000 to 0.300 in May 2021 with a mean and standard error of 0.064 ± 0.031. In May 2022, the single locus values ranged from 0.000 to 0.500 with a mean and standard error of 0.058 ± 0.049. The genetic diversity for five loci remained consistent among time points, while three loci slightly increased and one decreased. Differences in genetic diversity among inter-annual time points for each locus ranged from -0.191 to 0.111. At AL- CRC, *H_E_* values were the same in May 2022 and April 2023 (*H_E_* = 0.159). In May 2022, the single locus values ranged from 0.000 to 0.740 with a mean and standard error of 0.159 ± 0.071. In April 2023, the single locus values ranged from 0.000 to 0.640 with a mean and standard error of 0.159 ± 0.073. The genetic diversity remained consistent between time points for four loci, decreased for three loci, and increased for three loci. Differences in genetic diversity among inter-annual time points for each locus ranged from 0.000 to 0.238 (Figure 2, Table 3). Allelic richness was also low among all interannual time points, ranging from 1.00 (May 2021 and July 2023 at OH-MCC) to 1.49 (May 2022 at AL-CRC; Table 3).

All inter-annual values of pareto *β* were < 0.7. Pareto *β* values were 0.01 to 0.02 at OH- MCC, 0.10 to 0.16 at AL-YEC, and 0.49 to 0.66 at AL-CRC (Table 3). Multilocus linkage disequilibrium (*r̄_d_*) ranged from 0.03 to 1.00 among sites and exhibited a greater amount of variation among sites rather than between time points sampled at a site (Table 3). At OH-MCC, |D’| could not be calculated for May 2021 and July 2023 because all loci were fixed. There were 53 comparisons from May 2022. All except four pairs had a |D’| of 0.00. The remaining four range from 0.04 to 0.96 with a mean and standard error of 0.06 ± 0.03. At AL-YEC, there were 99 pairwise comparisons in May 2021, with a range of 0.00-0.94 and a mean and standard error of 0.08 ± 0.02. In May 2022, there were 64 pairwise comparisons with a range of 0.00-0.55 and a mean and standard error of 0.02 ± 0.01. At AL-CRC, there were 156 pairwise comparisons in May 2022 with a range of 0.00-0.85 and a mean and standard error of 0.09 ± 0.01. In April 2023, there were 152 pairwise comparisons with a range of 0.00-0.96 and a mean and standard error of 0.11 ± 0.02 (Figure 3, Table S5).

**Figure 3.**
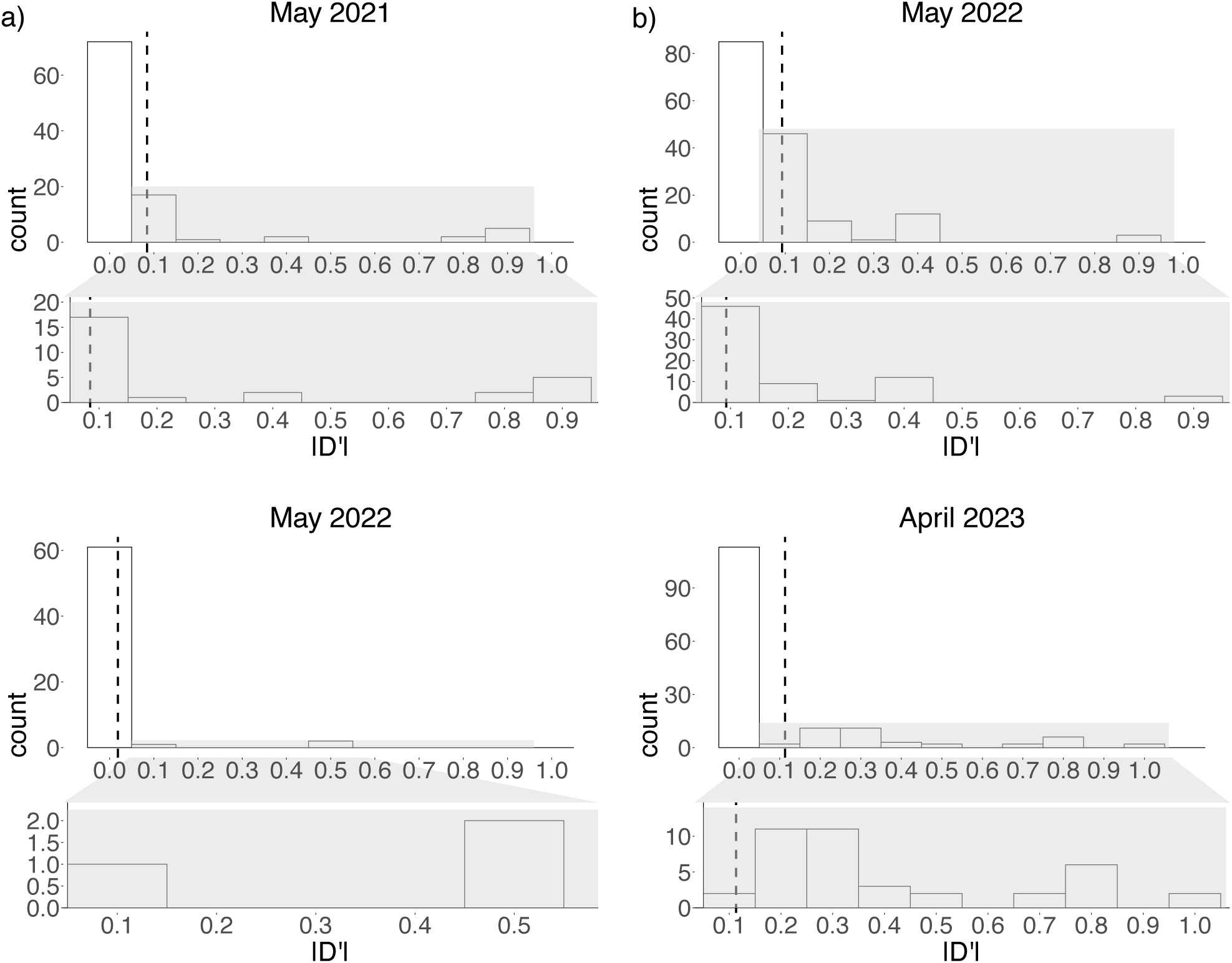
The discretized distribution of pairwise linkage disequilibrium (|D’|) values per locus are shown the for inter-annual time points in a) Yellow Creek (AL-YEC), and b) Cripple Creek (AL-CRC). The x-axis indicates discretized linkage disequilibrium values by 0.1 intervals from 0.0 to 1.0 and the y-axis (“count”) indicates the number of loci with a given |D’|range of values. Black dashed lines indicate the mean |D’|value. The |D’|value was 0.0 when both alleles in a pair were the same.

For each site, we could not calculate values of temporal genetic differentiation for most pairs of loci due to a lack of allelic variation. Those pairs that could be calculated were close to zero, suggesting little differentiation among inter-annual time points (Figure 4, Table S6).

**Figure 4.**
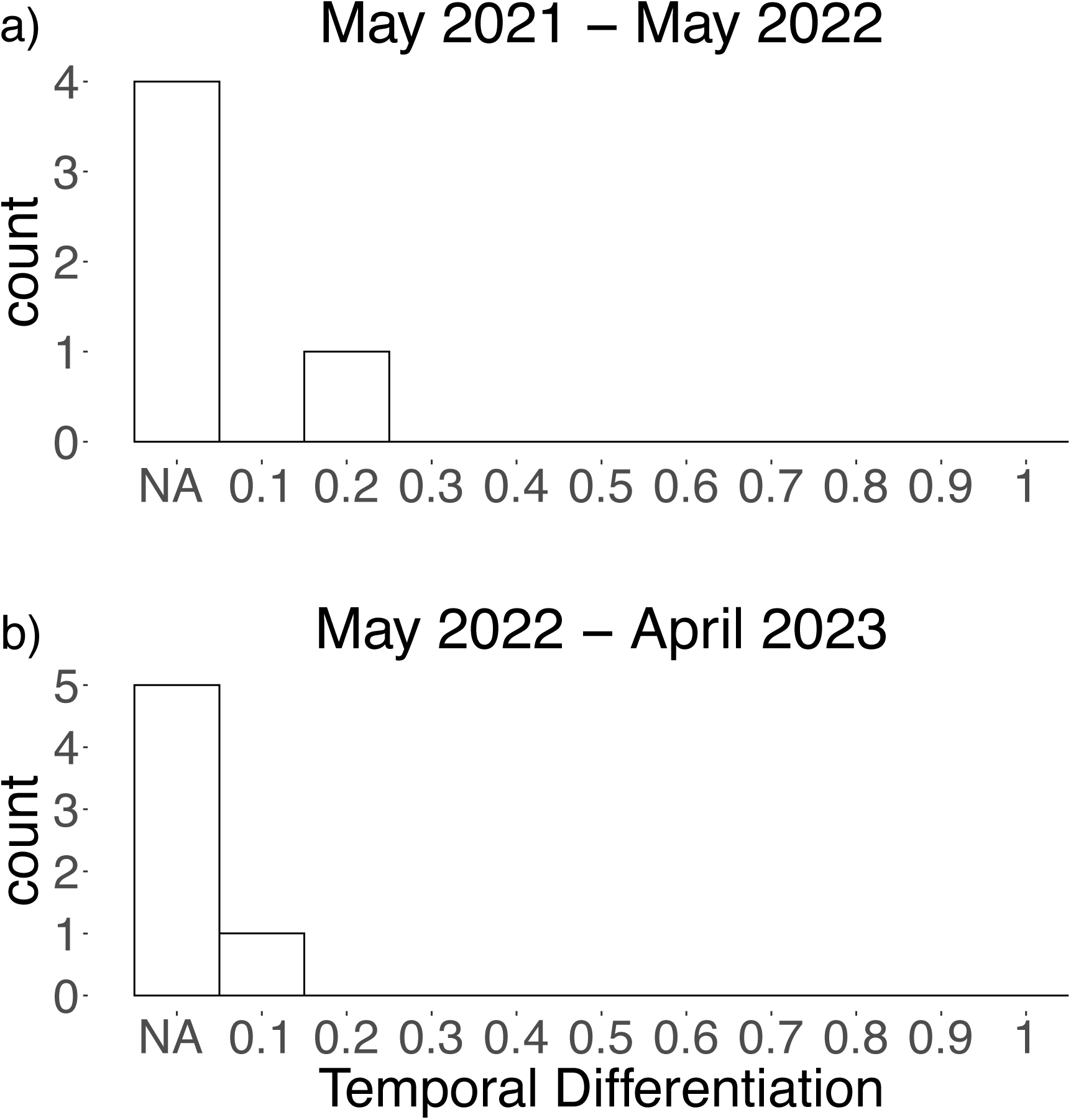
The distribution of temporal differentiation is shown for inter-annual time points in a) Yellow Creek (AL-YEC) and b) Cripple Creek (AL-CRC). The x-axis indicates temporal differentiation (measured as *F_ST_* between time points for each locus; ranges from 0-1) and the y- axis (“count”) indicates the number of loci with a given temporal differentiation range of values. If the expected heterozygosity (*H_E_*) for a locus at both time points was “0”, the pairwise temporal differentiation is indicated as “NA” – not applicable – as differentiation could not be calculated.

At AL-CRC, there were fewer fixed alleles compared to OH-MCC and AL-YEC (Table 3). At OH-MCC, there were no diverging alleles between most pairs, because all except two gametophytes from May 2022 belonged to the same MLG. At AL-CRC, the range (0 to 4) and mean (∼1.5) of diverging allele counts remained consistent at both time points. Additionally, at locus Bgel_071, there was an increase in the raw number of alleles from one year to the next (Table 3). At AL-YEC, the diverging alleles distribution ranged from 0 to 4 in May 2021 and from 0 to 1 in May 2022. However, the mean (∼0.5) remained consistent between time points because most pairs in May 2021 exhibited either 0 or 1 diverging alleles (Figure 5).

**Figure 5.**
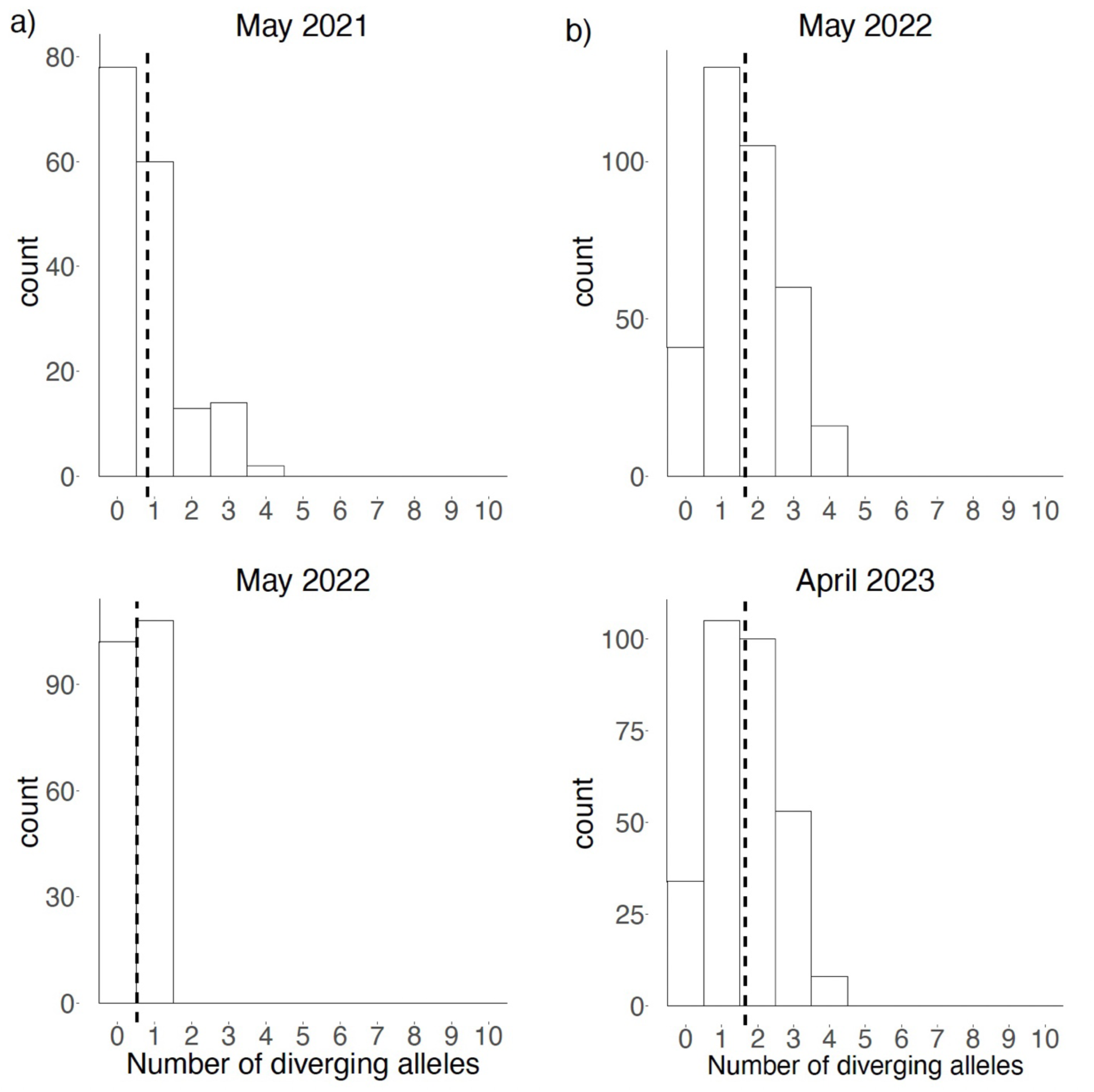
The distribution of counts of diverging alleles between each pair of gametophytes are shown for the inter-annual time points in a) Yellow Creek (AL-YEC) and b) Cripple Creek (AL- CRC). The x-axis represents the number of diverging alleles from 0 to 10. The y-axis (“count”) represents the number of pairs of gametophytes with the given number of diverging alleles. Black dashed lines indicate the mean number of diverging alleles.

### Intra-annual comparisons

The *pid* values remained consistent within intra-annual time points sampled at the same site. At MI-TRB, *pid* ranged from 2.17 x 10^-5^ in May 2022 (n=6) to 4.76 x 10^-5^ in July 2022 (n=13). At AL-CRC, *pid* ranged from 0.002 in March 2023 to 0.012 in April 2023 (n=25). At AL-CRC, *pid* ranged from 0.004 in May 2022 to 0.012 in April 2023 (Table 3).

Genotypic richness and evenness were stable between time points. At MI-TRB, genotypic richness (*R*) decreased slightly from 0.60 in May 2022 to 0.42 in July 2022, while *D\** also decreased slightly from 0.80 in May 2022 to 0.64 in July 2022 (Table 3). No MLGs were repeated between both time points. At AL-CRC, *R* decreased slightly from 0.54 in March 2023 to 0.42 in April 2023, while *D\** remained consistent at 0.89 at both time points (Table 3). Three MLGs were shared between time points, five unique MLGs were sampled only in March 2023, and four unique MLGs were sampled only in April 2023.

Site MI-TRB had slightly greater genetic diversity (*H_E_*) as compared to AL-CRC. Between intra-annual time points at a site, *H_E_* was stable and low (Table 3). At MI-TRB, single locus genetic diversity values ranged from 0.000 to 0.670 in May 2022, with a mean and standard error of 0.344 ± 0.070. In July 2022, single locus *H_E_* values ranged from 0.000 to 0.500 with a mean and standard error of 0.308 ± 0.057. Two loci remained consistent between time points, two decreased, and six increased (Figure 6). At AL-CRC, *H_E_* was 0.159 to 0.190 (Table 3).

**Figure 6.**
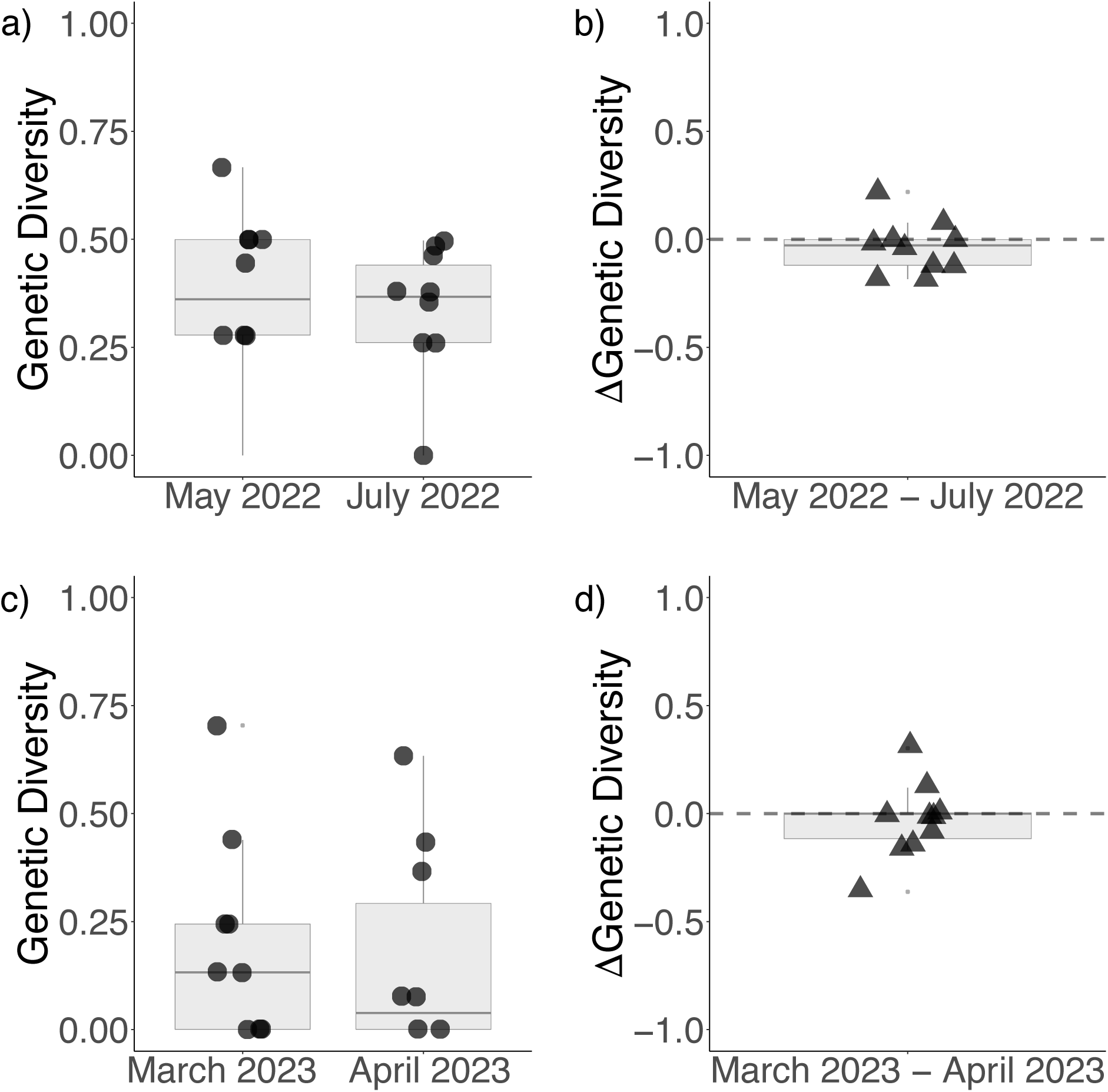
The distribution of genetic diversity (calculated as expected heterozygosity, *H_E_*) values at intra-annual time points are shown for a) Traverse River (MI-TRB) and c) Cripple Creek (AL- CRC). The y-axis range is shown as 0 to 1 for genetic diversity values in a) and c). Changes in genetic diversity within years are also indicated for b) Traverse River (MI-TRB) and d) Cripple Creek (AL-CRC). The y-axis range is shown as -1.0 to 1.0 for the change in genetic diversity estimates per locus in b) and d), with a dashed grey line to indicate the y-intercept at 0. Boxes represent the interquartile range, the middle lines are medians, whiskers represent the 1.5 interquartile ranges, and the small light grey dots represent outliers. Boxes represent the interquartile range, the middle lines are medians, whiskers represent the 1.5 interquartile ranges, and the small light grey dots represent outliers.

Single locus values ranged from 0.000 to 0.700 in March 2023 with a mean and standard error of 0.190 ± 0.073. In April 2023, single locus values ranged from 0.000-0.630 with a mean and standard error of 0.159 ± 0.073. Values for four loci remained consistent between time points, two decreased, and four increased (Figure 6). Site MI-TRB also had slightly higher mean allelic richness compared to AL-CRC, with values of 2.30 in May 2022 and 1.97 in July 2022; whereas AL-CRC had values of 1.60 in March 2023 and 1.42 in April 2023 (Table 3).

The pareto *β* values at both sites decreased between intra-annual time points: at MI-TRB, from 0.63 in May 2022 to 0.23 in July 2022, and at AL-CRC, from 0.84 in March 2023 to 0.66 in April 2023 (Table 3). Multilocus linkage disequilibrium (*r̄_d_*) values decreased slightly between intra-annual time points at both sites: from 0.66 to 0.50 at MI-TRB and from 0.20 to 0.04 at AL- CRC (Table 3). Pairwise values of linkage disequilibrium among alleles (|D’|) were also calculated. At MI-TRB, there were 234 pairwise comparisons for May 2022, with a range of 0.00 to 0.83 and a mean and standard error of 0.40 ± 0.02. There were 297 pairwise comparisons for July 2022, with a range of 0.00 to 0.92 and a mean and standard error of 0.36 ± 0.02. For most loci, |D’| decreased rather than increased through time. At AL-CRC, there were 173 pairwise comparisons in March 2023, with a range of 0.00 to 0.86 and a mean and standard error of 0.17 ± 0.02. For April 2023, there were 152 pairwise comparisons with a range of 0.00 to 0.96, and a mean and standard error of 0.11 ± 0.02. The mean |D’| value slightly decreased through time for both sites (Figure 7, Table S5).

**Figure 7.**
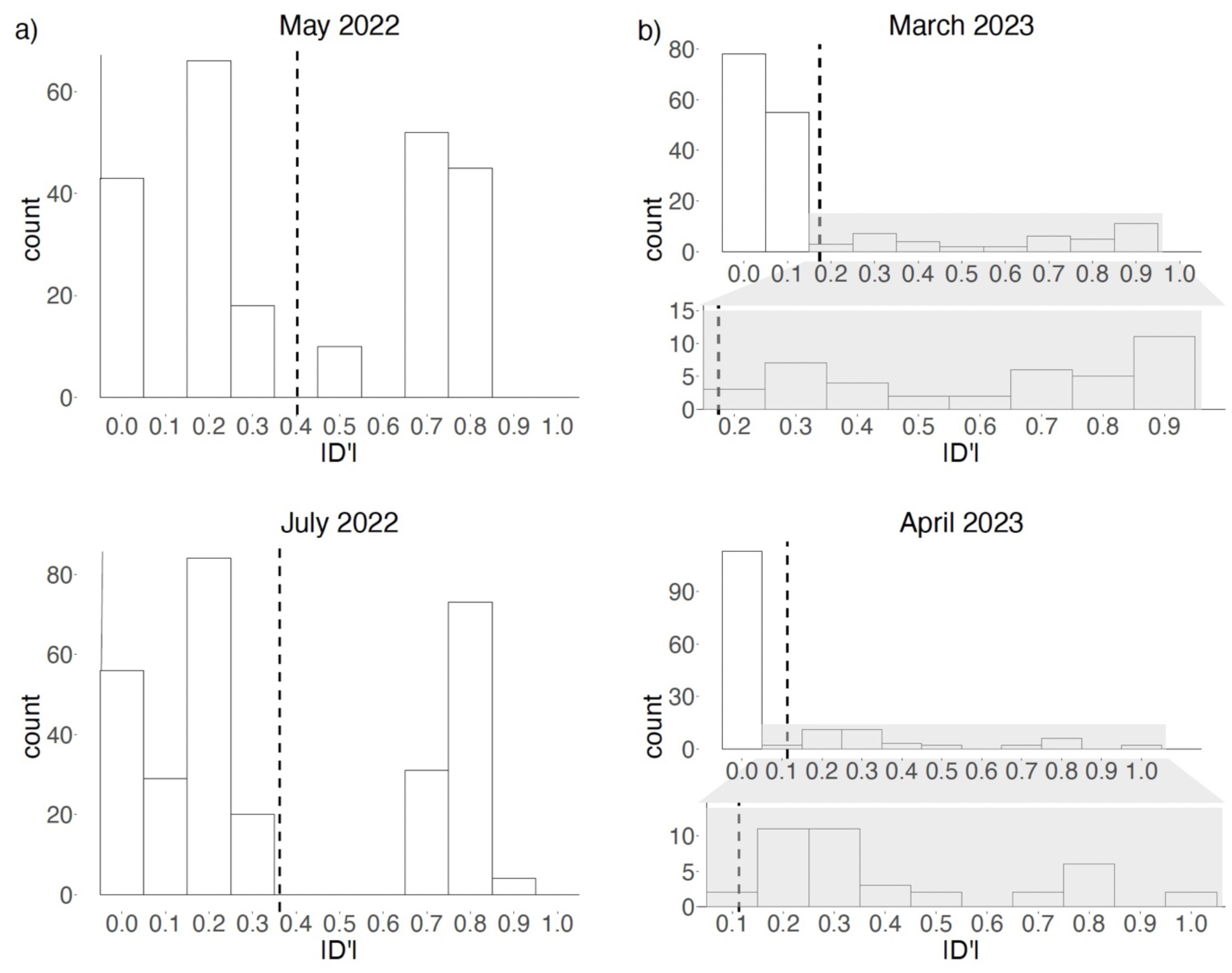
The discretized distribution of pairwise linkage disequilibrium (|D’|) values per locus are shown the for intra-annual time points in a) Traverse River (MI-TRB) and b) Cripple Creek (AL-CRC). The x-axis indicates per-locus linkage disequilibrium (|D’|, ranges from 0-1) and the y-axis (“count”) indicates the number of loci with a given |D’|value. Black dashed lines indicate the mean |D’| value.

We assessed genetic differentiation between intra-annual time points by calculating pairwise values for each locus. At AL-CRC, differentiation could not be calculated between most pairs of loci due to a lack of variation. At MI-TRB, more differentiation values could be calculated. Though genetic differentiation was relatively low between May 2022 to July 2022, values were slightly higher than those that could be calculated between intra-annual time points for AL-CRC and those that could be calculated for inter-annual time points at all sites (Figure 4, Figure 8, Table S6).

**Figure 8.**
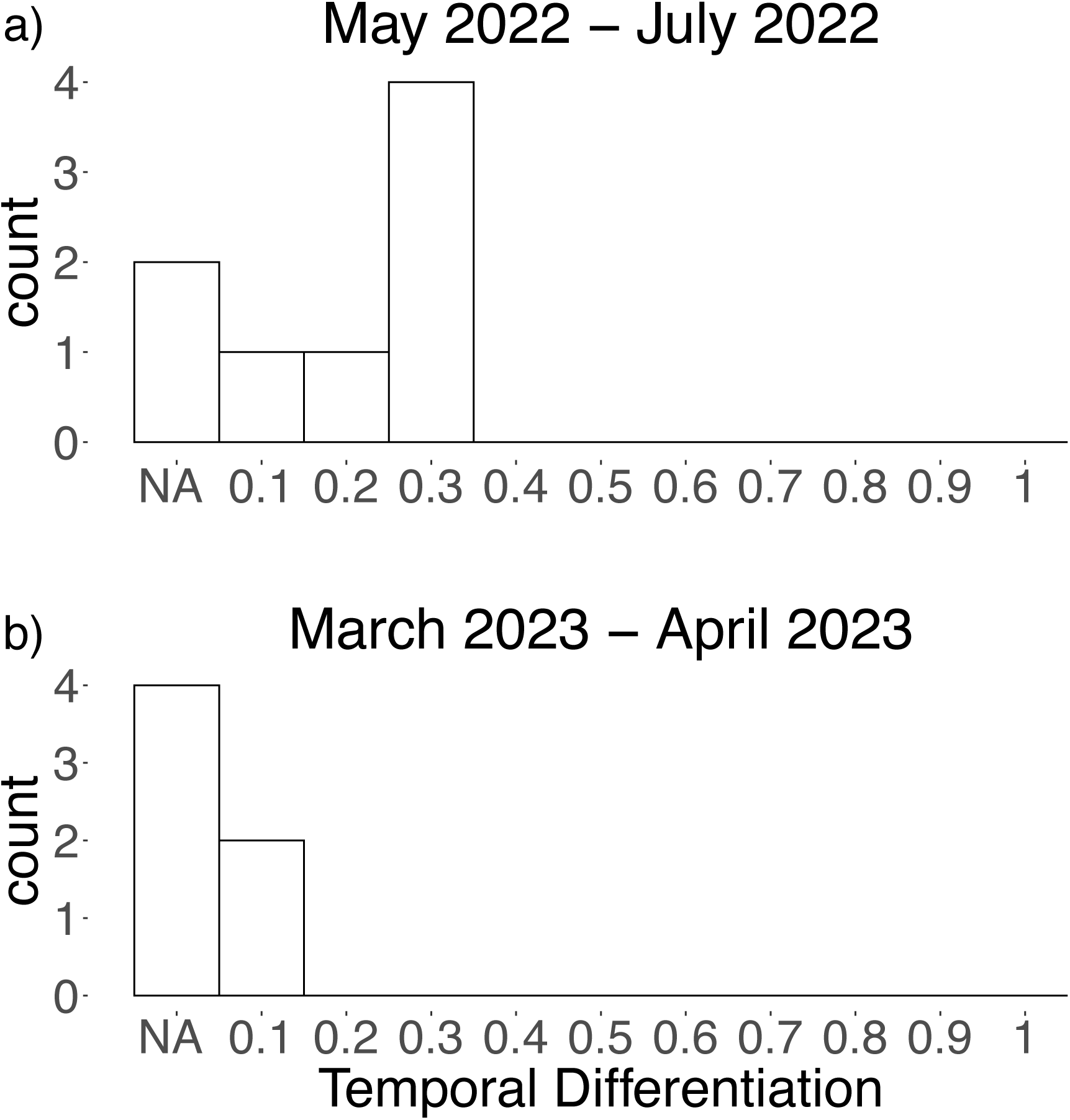
The distribution of temporal differentiation is shown for intra-annual time points in a) Traverse River (MI-TRB) and b) Cripple Creek (AL-CRC). The x-axis indicates temporal differentiation (measured as *F_ST_* between time points for each locus; ranges from 0-1) and the y- axis (“count”) indicates the number of loci with a given temporal differentiation range of values. If the expected heterozygosity (*H_E_*) for a locus at both time points was “0”, the pairwise temporal differentiation is indicated as “NA” – not applicable – as differentiation could not be calculated.

The number of fixed alleles remained consistent between intra-annual time points sampled at a site. Site MI-TRB had fewer fixed alleles than AL-CRC (Table 3). At both sites, the mean and range of the number of diverging alleles remained consistent between time points, with slight shifts in the distributions. The diverging allele counts implied more genetic variation at MI-TRB than at AL-CRC, but with similar patterns between intra-annual time points within each site (Figure 9).

**Figure 9.**
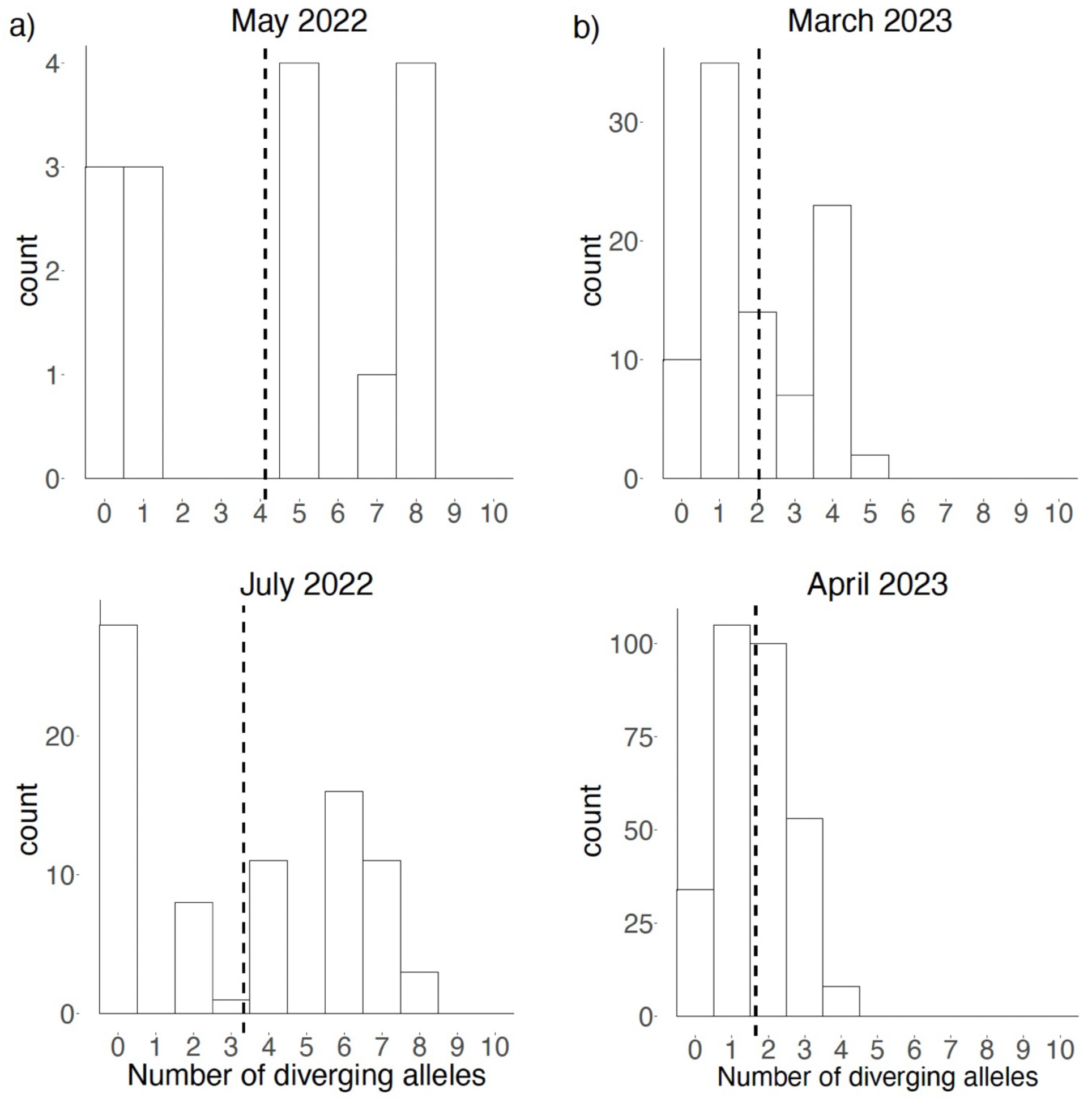
The distribution of counts of diverging alleles between each pair of gametophytes collected at intra-annual time points in a) Traverse River (MI-TRB) and b) Cripple Creek (AL- CRC). The x-axis represents the number of diverging alleles. The y-axis (“count”) represents the number of pairs of gametophytes with the given number of diverging alleles. Black dashed lines indicate the mean number of diverging alleles.

## DISCUSSION

We measured temporal variation in genetic diversity to better assess the reproductive system of *Batrachospermum gelatinosum*. The *pid* values for sites genotyped with ten loci indicate that we have sufficient resolution of genotypes, but this was not the case for OH-MCC (Ohio). This could be due to high rates of intragametophytic selfing at this site, leading to consistently low diversity. At MI-TRB (Michigan), we surmise there are higher outcrossing rates than the other sites based on greater diversity. Both AL-CRC and AL-YEC (Alabama) were intermediate between MI-TRB and OH-MCC (see also Shainker-Connelly et al., 2024). There was little temporal partitioning of genetic variation. Below, we discuss how inter-annual comparisons inform our knowledge of the reproductive system and how intra-annual comparisons inform our understanding of the seasonality of the *B. gelatinosum* life cycle.

### Inter-annual comparisons: implications for the reproductive system

At the sites we sampled, the reproductive system of *Batrachospermum gelatinosum* seems to remain consistent with low variation in genetic and genotypic diversity among time points. Low genetic diversity is likely maintained by high rates of intragametophytic selfing, monospore production, or both. We interpret these data as intragametophytic selfing rather than monospore production because we collected gametophytes from all our sampling sites, including those with low genotypic diversity. The presence of gametophytes indicates that meiosis has occurred as gametophytes are unlikely to result from any clonal process. Moreover, we have observed very small gametophytes (∼1 cm) bearing many carposporophytes (Shainker-Connelly, Crowell, Vis, and Krueger-Hadfield, personal observations). Therefore, fertilization is likely efficient and occurs when gametophytes are very small. Nevertheless, the rates of monospore production by chantransia in natural populations need to be quantified to determine the extent to which clonality contributes to the patterns we observed.

The results of inter-annual comparisons at site OH-MCC are consistent with our predictions for low standing levels of genetic diversity associated with high rates of intragametophytic selfing (Table 1). As we sampled one dominant genotype, it is likely that intragametophytic selfing has been maintained long-term at this site. The results at site AL-YEC are also consistent with high rates of intragametophytic selfing, though there is likely a slightly higher level of standing genetic diversity than at OH-MCC (Table 1). While the pareto *β* values indicate intragametophytic selfing, the multilocus and single locus values of linkage disequilibrium are both lower at AL-YEC than at OH-MCC. Additionally, the slight decrease in the mean number of diverging alleles from 2021 to 2022 at AL-YEC may have resulted from intragametophytic selfing eroding genetic diversity from only one generation to the next.

However, it is also possible that the slight differences observed could be due to chance. It would be necessary to sample additional time points with larger sample sizes, if possible, to determine whether this pattern reflects natural processes or is the result of sampling error. An alternative explanation is that instead of a change in the reproductive system through time, the chantransia that were the product of intragametophytic selfing or monospore production from previous years later produced gametophytes. Future studies would need to genotype chantransia through time to test the feasibility of this explanation.

The results from AL-CRC gametophytes are consistent with our predictions for greater standing genetic diversity with a mixed reproductive system that may include higher rates of outcrossing compared to the other sites. This prediction is supported by low *pid* values compared to OH-MCC and AL-YEC, greater pareto *β* values, and a greater mean and wider distribution of the number of diverging alleles. The mixed reproductive system at this site may be driven by greater heterogeneity in biotic and abiotic factors, as a mixed mating system can be stable if there is temporal variation in resource availability (Bengtsson & Ceplitis, 2000; Schemske & Lande, 1985; Weeks, 1993) and environmental fluctuations likely drive reproductive system evolution (Pierre et al., 2022). The microhabitats available at this site are more heterogeneous than other sites we sampled in this study. There were two riffles and a pool present within the sampling area and gametophytes were collected from all these microhabitats, though they were less abundant in the pool than the riffles. The reproductive mode varies by microhabitat in other taxa (e.g., water availability, see Johnson & Shaw, 2015), so this variability may contribute to the patterns observed at AL-CRC. For example, the higher flow velocity in riffles may facilitate greater fertilization rates than the lower flow velocity in pools. This site was also very diverse in terms of the number of freshwater red algal species observed (Table S7), and it is possible that there is a relationship between the species and genetic diversity (Vellend & Geber, 2005).

Similar selective factors could drive diversification both within and among species, or species diversity may influence the selection regime that drives genetic diversity (Vellend & Geber, 2005). For example, Van Valen (1965) hypothesized that in species-rich communities, the niche breadth of each species would be smaller due to competition, driving a decrease in genetic diversity. Conversely, Harper (1977) predicted that species diversity acts as a source of diversifying selection, so that there is a positive relationship between species and genetic diversity.

Though there were slight differences among sites, the reproductive system likely remains stable through time. Genotyping these sites in future years would be useful to distinguish the time scales at which natural selection may act on the traits that affect the reproductive system.

Genotyping chantransia would enable us to measure heterozygosity and quantify clonal rates. Additionally, future studies should investigate sites that likely have lower rates of uniparental reproduction than those included here (e.g., site MI-CUT in Shainker-Connelly et al., 2024). Including sites with lower rates of uniparental reproduction and across more geographic areas would be useful to determine whether stability of the reproductive system is a universal pattern across geographically and genetically distinct sites.

### Intra-annual comparisons: Implications for the seasonality of the life cycle

We characterized intra-annual temporal patterns at two sites – MI-TRB and AL-CRC. At both sites, the first sampling point was early in the season when gametophytes were small and patchily distributed throughout the sampled reach. The second sampling point was closer to the peak of gametophytic abundance, when gametophytes were larger and more continuously distributed throughout the sampled reach. Both sites exhibited a slight decrease in linkage disequilibrium between time points, which may be associated with the increase in population size (Waples, 2006), but alternatively may be artifacts of sample sizes at the earlier sampling points.

At MI-TRB, no MLGs were shared between time points. In May 2022, the gametophytic subpopulation was small and patchy. Only eight gametophytes were collected and genotyped, from which two were removed because of non-amplification at one locus each. Consequently, it is possible that the sample size at the May time point did not sufficiently capture the genotypes present or possible following meiosis. However, several measures, including allelic richness, genetic diversity, and genotypic richness, suggest that this site is more genetically and genotypically diverse compared to most other *Batrachospermum gelatinosum* sites (see also Shainker-Connelly et al., 2024). Patterns observed at site MI-TRB were consistent with the prediction that different chantransia genotypes may produce gametophytes at different times, possibly due to environmental changes through the season. A similar pattern has been observed in microalgal blooms, where genetic changes within seasons are thought to be driven by the germination of resting cysts under different conditions (i.e., variations in temperature; Lebret et al., 2012). Alternatively, there could be a succession of genotypes either adapted to slightly different environmental conditions or stochastically favored by disturbance events throughout the season, as observed in rotifers for which a succession of different clonal groups corresponded with temporal changes in environmental conditions (Gómez et al., 1995).

The population genetic diversity and structure of AL-CRC remained more consistent compared to MI-TRB, with little differentiation between intra-annual time points. Several genotypes were found at just one time point and not the other, but we did not observe turnover of MLGs to the same degree as at MI-TRB. It should be noted that the intra-annual time points at AL-CRC were sampled just one month apart, while MI-TRB was sampled two months apart.

More genotypic shifts may also occur at AL-CRC if observed over a longer two-month period; however, in 2023, gametophytes at AL-CRC were no longer present two months after our initial time point.

Overall, these results suggest that when making population genetic comparisons between and among sites, it may be important to consider the month of sampling and the length of time in which gametophytes are present. Environmental changes and disturbances throughout the season, such as snow melt in northern sites like MI-TRB (Michigan) and storms with heavy rainfall in southern sites like AL-CRC (Alabama), may drive the genotypic shifts observed. Future studies should aim to sample at a finer time scale and to measure both population size and environmental variables to better understand the environmental, genetic, and demographic dynamics of gametophytes through time.

### Drivers of temporal genetic structure

Temporal environmental changes can cause shifts in the reproductive system which in turn shape population genetic structure (Eckert et al., 2010). As habitat disturbance and fragmentation increase, plants tend to shift towards higher rates of inbreeding and more variable outcrossing rates (Coates et al., 2013; Eckert et al., 2010). Additionally, it is likely advantageous for both sexual and asexual reproduction to be maintained when conditions are environmentally variable, and the proportion of each type of reproduction may vary temporally as environmental conditions change (Bengtsson & Ceplitis, 2000). For example, both microalgal blooms (Lebret et al., 2012) and cyclical parthenogens, such as potato-peach aphids (Guillemaud et al., 2003), have mixed reproductive modes in which asexual reproduction occurs throughout the year and sexual reproduction may occur once per year. In these situations, there are intra-annual shifts in population genetic structure driven by cyclical reproductive modes, but a stable mixed reproductive mode is maintained long-term (Lebret et al., 2012; Guillemaud et al., 2003). It is possible that a similar pattern occurs in freshwater red algae. A stable mixed reproductive mode may be maintained where the chantransia reproduce asexually via monospore production throughout the year, meiosis occurs during one season to produce gametophytes, and then fertilization only occurs during the season in which gametophytes are present. Future studies could assess the niche differences between chantransia and gametophytes to determine whether monospore production is favored by certain environmental conditions.

Although there was little inter-annual variation in our study, there were more differences in genotypes between intra-annual time points, especially at MI-TRB. This pattern is more likely driven by the seasonality of the life cycle rather than the reproductive system. Chantransia are likely present throughout the year, but gametophytes are typically present only during a specific season. These life cycle shifts may be driven by the environment. For example, gametophyte cover may be positively correlated with stream depth, current velocity, and day length (Drerup & Vis, 2014), as well as substratum type (Hambrook & Sheath, 1991; Higa et al., 2007); and the timing of gametophytic reproductive maturity may be influenced by light availability and current velocity (Filkin & Vis, 2004). Additionally, chantransia may act as a repository for genetic diversity between seasons. In the bloom-forming microalga *Gonyostomum semen*, fertilization results in resting cysts that lie dormant in sediment through the winter (Figueroa & Rengefors, 2006). In the spring, these cysts germinate, and differences in environmental conditions may select for the germination of different genotypes (Lebret et al., 2012). Chantransia may act as a similar repository of genetic diversity by providing a persistent “bank of microscopic forms” (Chapman, 1987; Hoffmann & Santelices, 1991; Schoenrock et al., 2020). It is possible that (i) different chantransia genotypes undergo meiosis to produce gametophytes under different environmental conditions, and (ii) different gametophytic genotypes reach reproductive maturity under different environmental conditions. To better understand the putative role of the chantransia phase as a repository for genetic diversity, future studies should determine the lifespan of this phase (i.e., do chantransia thalli persist through multiple gametophyte seasons), the rate of reproduction via monospores, and the environmental factors that may trigger chantransia to undergo meiosis to produce gametophytes.

### Methodological limitations and future directions

Our data provided an overview of temporal patterns of population genetic diversity and structure and resulted in identifying future avenues to better understand this freshwater red macroalga. However, the population genetic tools needed to directly quantify clonal and sexual rates from the haploid phase of haploid-diploid organisms have not yet been developed. In fact, there is generally a lack of standardization in studies that assess clonal rates (Arnaud-Haond et al., 2007). Previous work (e.g., Lebret et al., 2012; Reynolds et al., 2017) relied on traditional genetic summary statistics, such as genetic differentiation (*F_ST_* measured between time points rather than between sites), genotypic diversity (*R*), inbreeding coefficient (*F_IS_*) and linkage disequilibrium (*r̄_d_*), to characterize the reproductive system. These statistics may be used as a proxy for clonal rates, but they are inaccurate for low to moderate clonal rates and do not disentangle the effects of intragametophytic selfing from asexual processes (Stoeckel et al., 2021; Shainker-Connelly et al., 2024). Additionally, *F_IS_* can only be calculated in the diploid phase. Here, we built upon previous studies by incorporating single locus values of these indices and considering their variance, which can be indicative of partial clonality (Stoeckel et al., 2021). We also incorporated measures of pareto *β* which can be used as a proxy for asexual reproduction and/or intragametophytic selfing (Arnaud-Haond et al., 2007). Future studies could distinguish between selfing and monospore production by using paternity analyses (e.g., Engel et al., 1999; Krueger-Hadfield et al., 2015) to directly assess rates of intra- and inter-gametophytic selfing. This is difficult with traditional methods due to the diminutive size of the carposporophytes in this taxon but is becoming more accessible as advancements in molecular biology will allow for the affordable genotyping of single cells and very small amounts of tissue (e.g., Börgstrom et al., 2017; Bowers et al., 2022).

Temporal genotyping may improve assessments of the reproductive system in partially clonal organisms. Becheler et al. (2017) developed methods to directly estimate clonal rates by calculating genotypic transitions between two separate time steps, but at present these methods are only suitable for diploid genotypes. Therefore, without modifying the life cycle model in this method, it would be necessary to sample and genotype the chantransia. Sampling the chantransia phase and performing single cell genotyping would allow for directly estimating clonal rates and the proportion of diploid chantransia to haploid gametophytes in the population. The proportion of haploids has profound consequences for the distribution of population genetic indices and calculation of clonal rates (Stoeckel et al., 2021). However, it is still difficult to collect morphological chantransia ‘individuals’ because they are microscopic. Developing new methods to directly assess clonal rates from the temporal sampling of haploids may thus be more pragmatic for *Batrachospermum gelatinosum*. Developing such methods, as recently achieved for polyploids (e.g., Stoeckel et al., 2024), will allow us to examine both spatial and temporal population genetic patterns across organisms with diverse life cycle types.

Our results suggest that the reproductive system remains consistent through short time scales and that the life cycle of *Batrachospermum gelatinosum* may drive genetic shifts within a season. The development of novel analytical tools, paired with finer scale temporal sampling, will improve our characterization of how the reproductive system and life cycle drive population genetic patterns. Applying these tools to other freshwater red algal taxa with varying sexual systems (see Krueger-Hadfield et al., 2024) will help us to determine whether patterns are consistent among freshwater reds. Expanding temporal reproductive system studies to include organisms with a wider variety of life cycle types will improve our understanding of the evolution of sex across eukaryotes.

## Acknowledgements

We thank M. Amsler, B. Anderson, W. Chiasson, A. Oetterer, B. Thornton, S. Thornton, and T. Williams for help collecting *B. gelatinosum*; A. Cao and M. Crowley for use of the capillary sequencer at the Heflin Center for Genomic Sciences at the University of Alabama at Birmingham (UAB); and C. Amsler, M. Sandel, and K. Marion for serving on the dissertation committee of SJSC and providing comments and suggestions that helped improve this manuscript. This manuscript represents the partial fulfillment of the PhD dissertation of SJSC.

## Data, scripts, code, and supplementary information availability

Will be uploaded to Zenodo

---

**Table S1.**
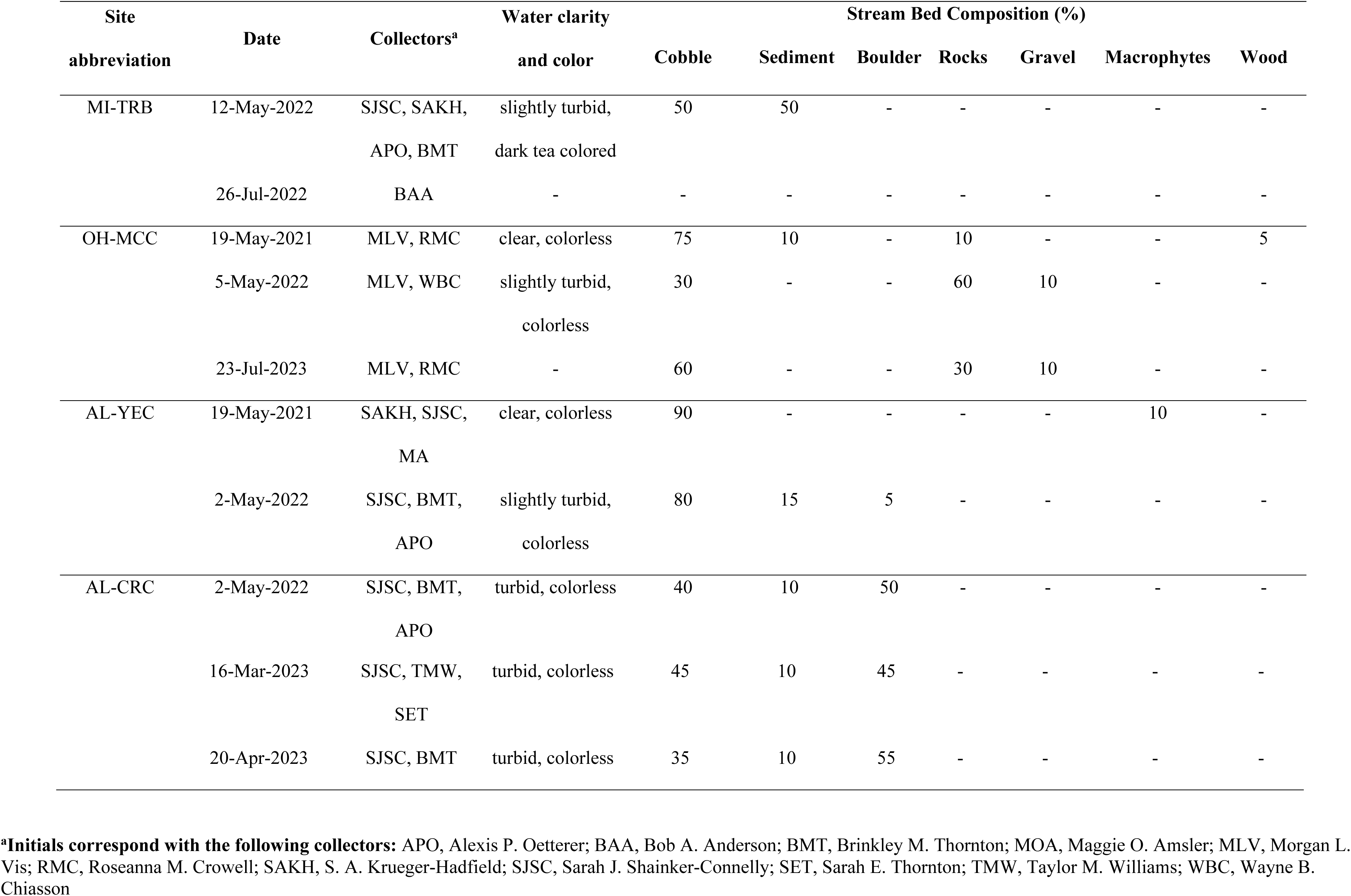
Water clarity and color, and stream bed composition for each date sampled per site. Site abbreviations as in Table 1.

| Site<br>abbreviation | Date | Collectors<sup>a</sup> | Water clarity | Stream Bed Composition (%) |  |  |  |  |  |  |
| --- | --- | --- | --- | --- | --- | --- | --- | --- | --- | --- |
|  |  |  | and color | Cobble | Sediment | Boulder | Rocks | Gravel | Macrophytes | Wood |
| MI-TRB | 12-May-2022 | SJSC, SAKH, | slightly turbid, | 50 | 50 | - | - | - | - | - |
|  |  | APO, BMT | dark tea colored |  |  |  |  |  |  |  |
|  | 26-Jul-2022 | BAA | - | - | - | - | - | - | - | - |
| OH-MCC | 19-May-2021 | MLV, RMC | clear, colorless | 75 | 10 | - | 10 | - | - | 5 |
|  | 5-May-2022 | MLV, WBC | slightly turbid,<br>colorless | 30 | - | - | 60 | 10 | - | - |
|  | 23-Jul-2023 | MLV, RMC | - | 60 | - | - | 30 | 10 | - | - |
| AL-YEC | 19-May-2021 | SAKH, SJSC,<br>MA | clear, colorless | 90 | - | - | - | - | 10 | - |
|  | 2-May-2022 | SJSC, BMT, | slightly turbid, | 80 | 15 | 5 | - | - | - | - |
|  |  | APO | colorless |  |  |  |  |  |  |  |
| AL-CRC | 2-May-2022 | SJSC, BMT, | turbid, colorless | 40 | 10 | 50 | - | - | - | - |
|  |  | APO |  |  |  |  |  |  |  |  |

|  |  |  |  |  |  |  |  |  |  |
| --- | --- | --- | --- | --- | --- | --- | --- | --- | --- |
| 16-Mar-2023 | SJSC, TMW, | turbid, colorless | 45 | 10 | 45 | - | - | - | - |
|  | SET |  |  |  |  |  |  |  |  |
| 20-Apr-2023 | SJSC, BMT | turbid, colorless | 35 | 10 | 55 | - | - | - | - |

**<sup>a</sup>Initials correspond with the following collectors:** APO, Alexis P. Oetterer; BAA, Bob A. Anderson; BMT, Brinkley M. Thornton; MOA, Maggie O. Amsler; MLV, Morgan L. Vis; RMC, Roseanna M. Crowell; SAKH, S. A. Krueger-Hadfield; SJSC, Sarah J. Shinker-Connelly; SET, Sarah E. Thornton; TMW, Taylor M. Williams; WBC, Wayne B. Chiasson

**Table S2.**
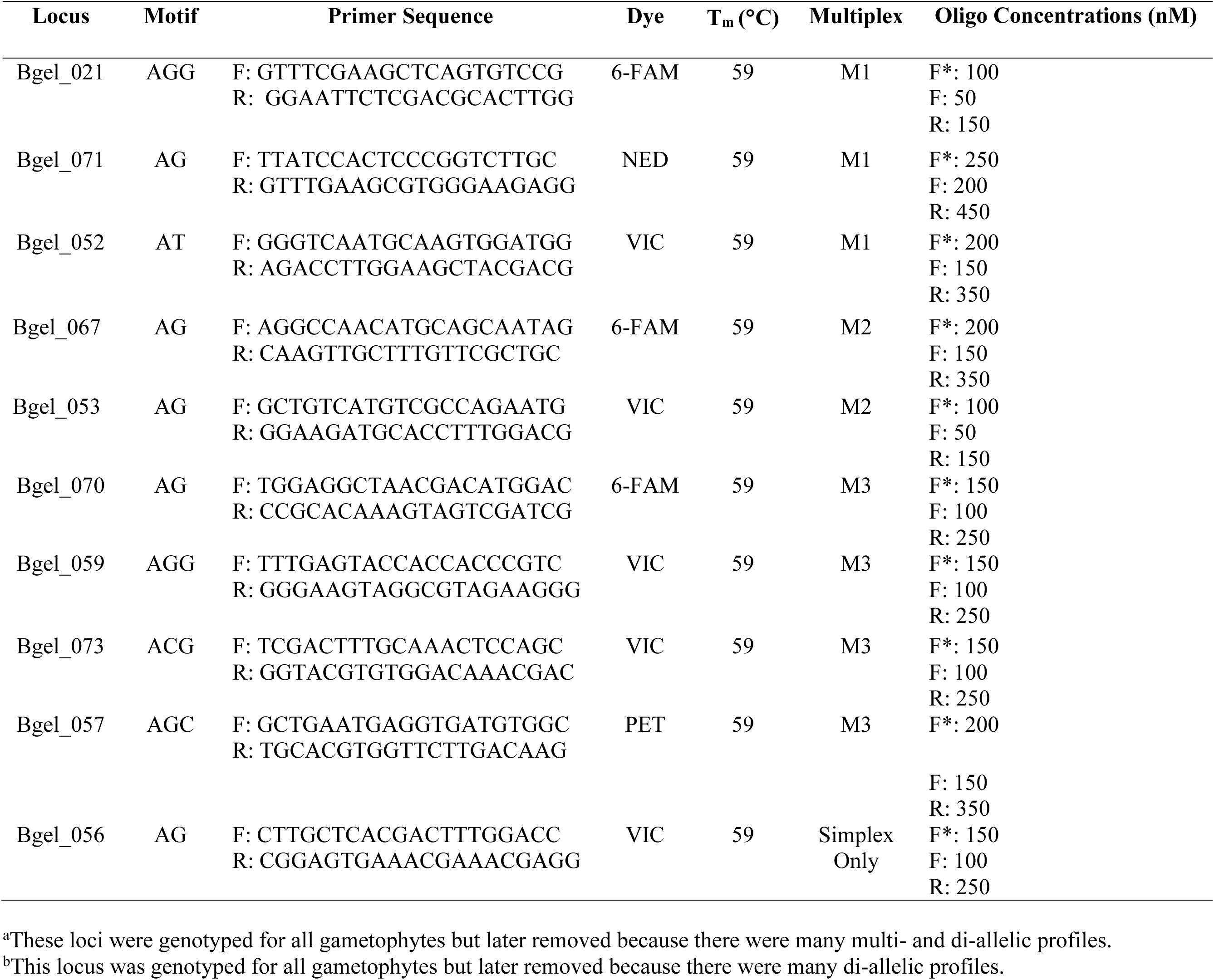
Oligo information for microsatellite loci used in genotyping *Batrachospermum gelatinosum* gametophytes. The assigned locus name, repeat motif, primer sequence, fluorochrome for the forward oligo, annealing temperature, and multiplex are given for all loci. Bgel_056 was always amplified in simplex PCR. Primer concentrations (nM) are given for the labeled forward (F*), unlabeled forward (F), and the unlabeled reverse (R) oligos.

**Table S3.**
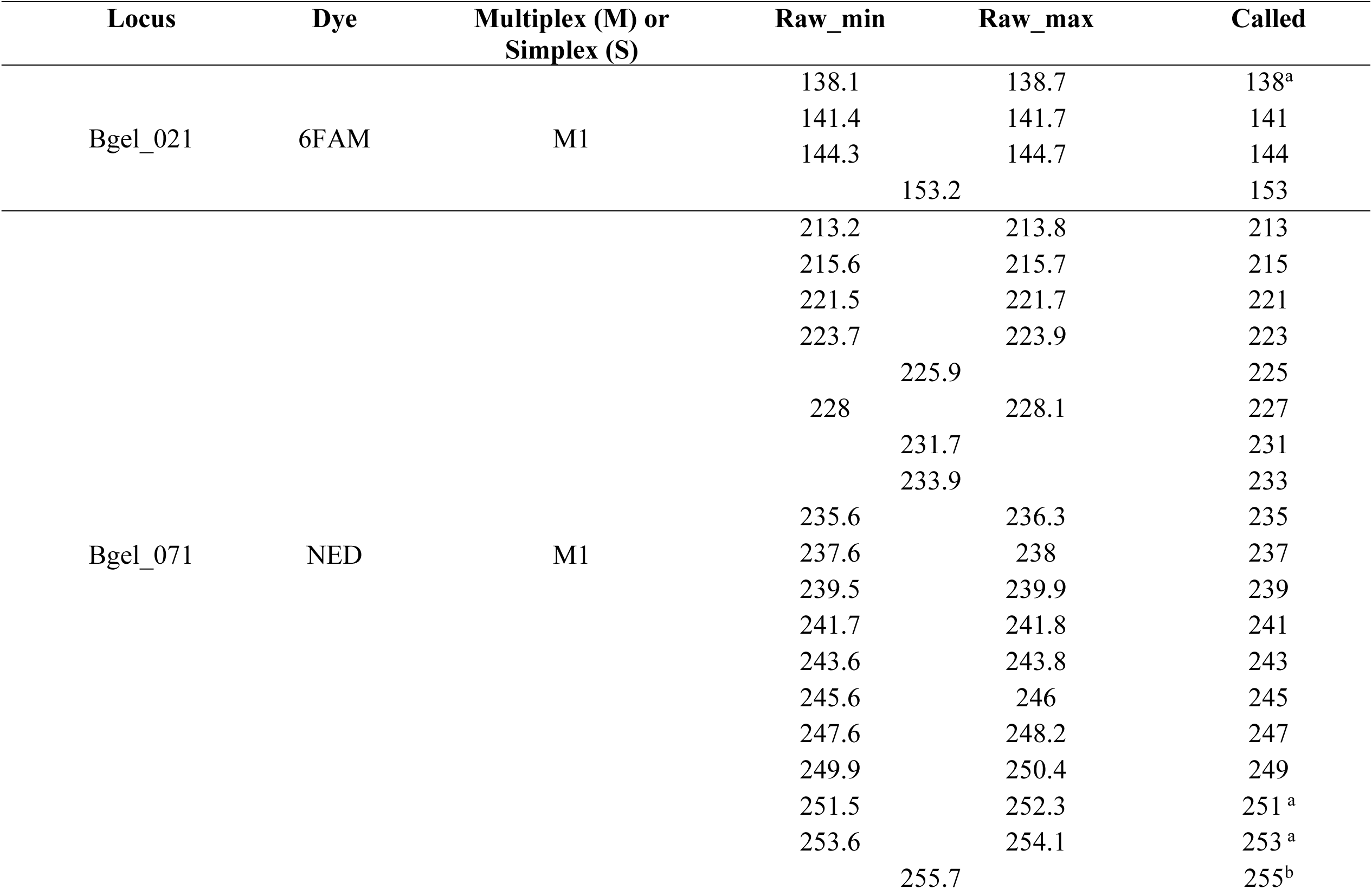
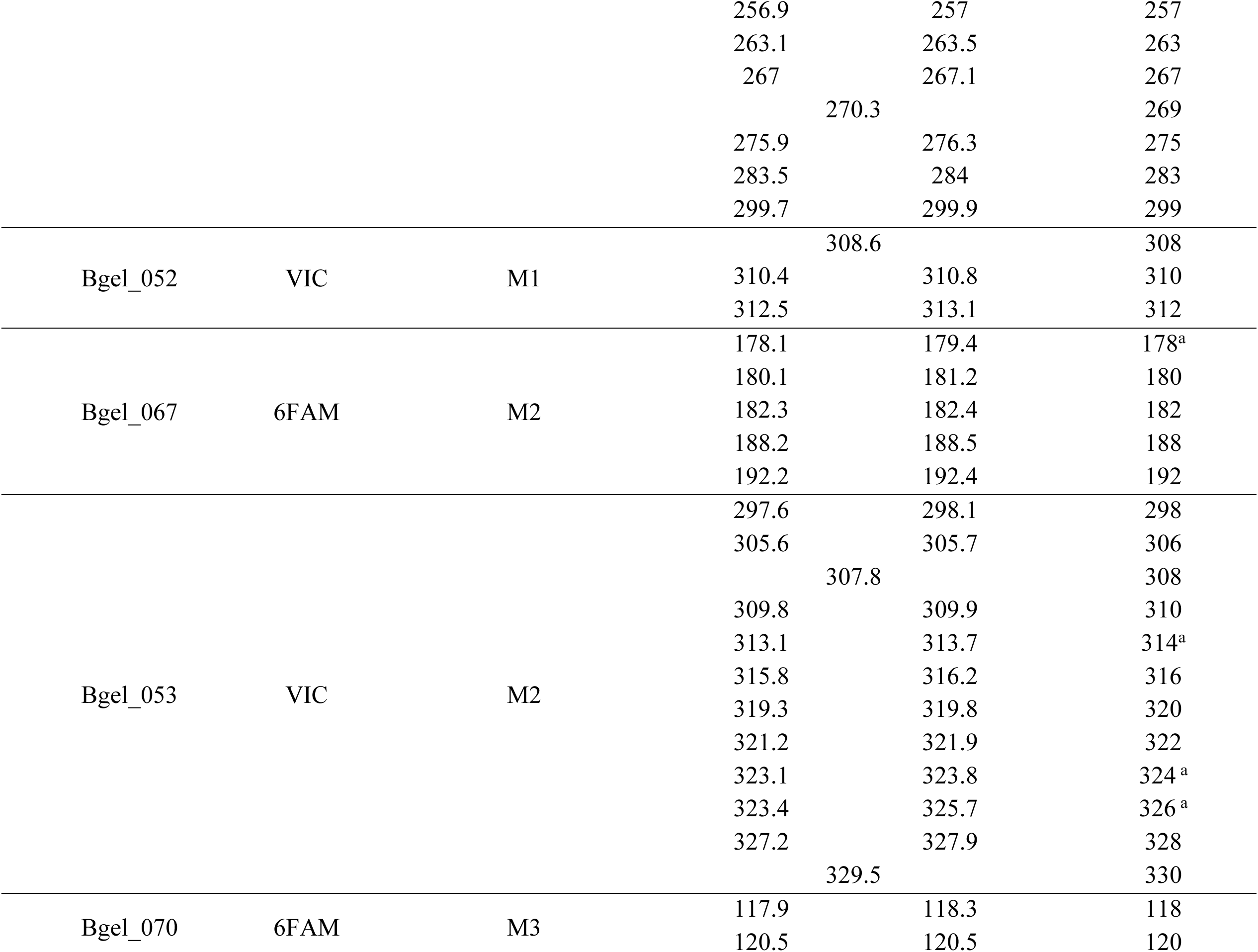
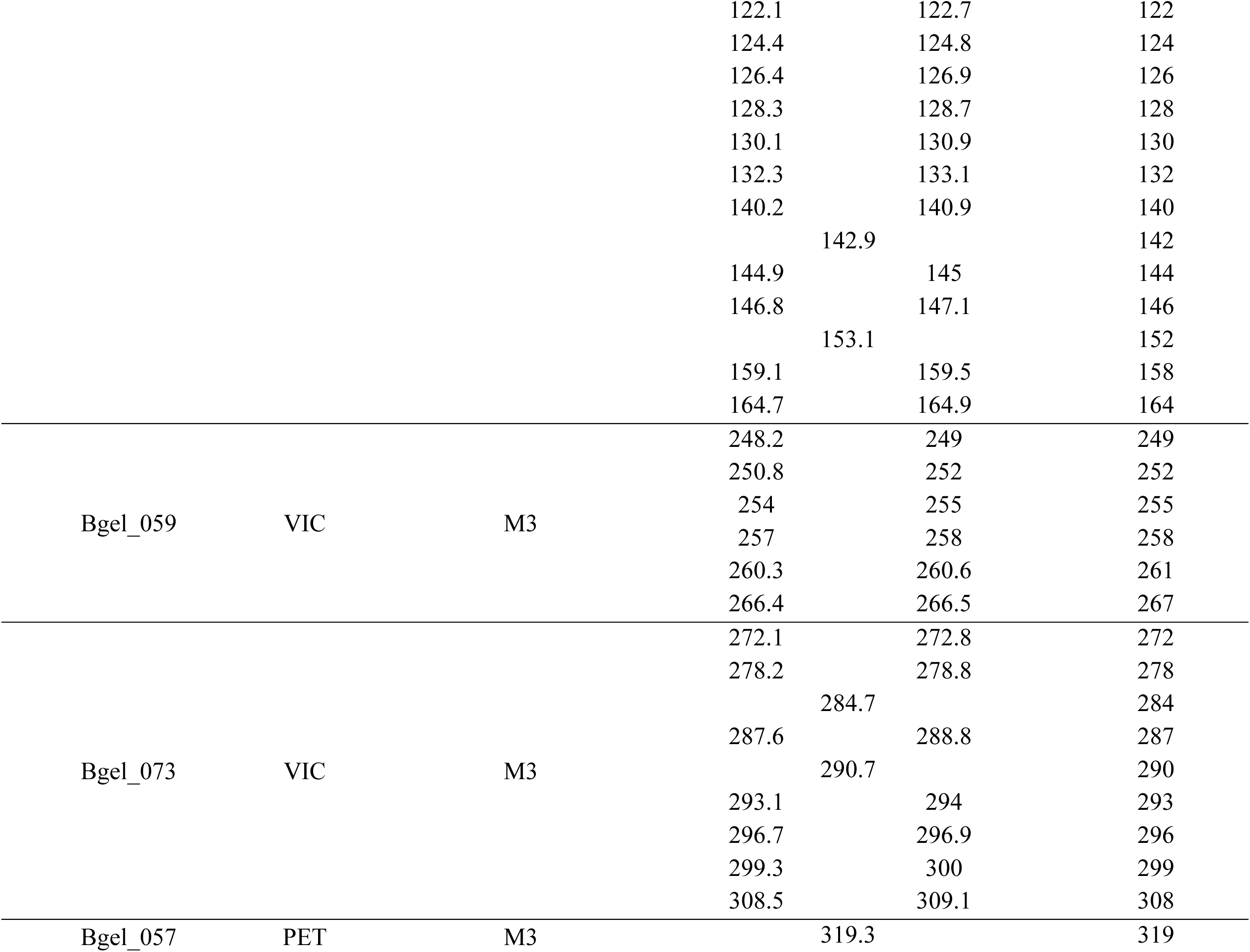
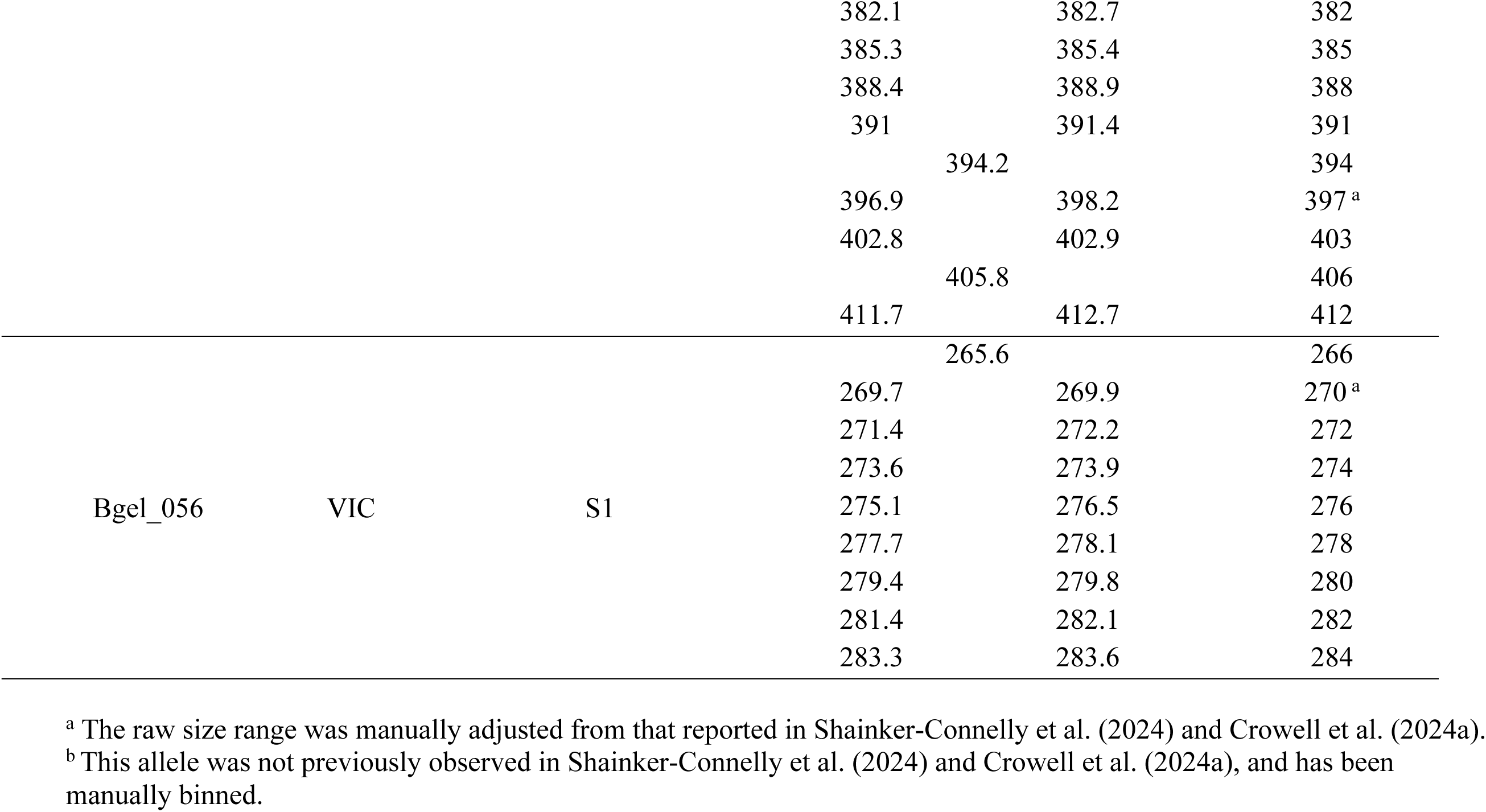
Raw size ranges for each binned allele. Each locus is given along with its fluorochrome and multiplex or simplex assignment. All alleles present for each locus are listed. The minimum and maximum raw allele sizes observed are given, along with the binned allele calls. If there was only one raw size for an allele, this size is listed between the raw_min and raw_max columns.

**Table S4.** Null allele frequencies for each locus were determined by non-amplification after 2-3 PCR attempts. As gametophytes are haploid, non-amplification of an allele at a given locus was considered a null allele (see also Krueger-Hadfield et al., 2013). Gametophytes with a null allele at any locus were removed for subsequent analyses.

| Locus | Null Allele Frequency |
| --- | --- |
| Bgel_021 | 0.5% |
| Bgel_071 | 31.3% (0.0%)<sup>a</sup> |
| Bgel_052 | 0.0% |
| Bgel_067 | 2.4% |
| Bgel_053 | 0.5% |
| Bgel_070 | 0.0% |
| Bgel_059 | 1.9% |
| Bgel_073 | 0.0% |
| Bgel_057 | 4.8% |
| Bgel_056 | 0.0% |

<sup>a</sup>Bgel\_071 did not amplify for most gametophytes from OH-MCC. The first null allele frequency includes the non-amplification of gametophytes from OH-MCC. The frequency in parentheses indicates the null allele frequency if OH-MCC is not included in the calculation.

**Table S5.**
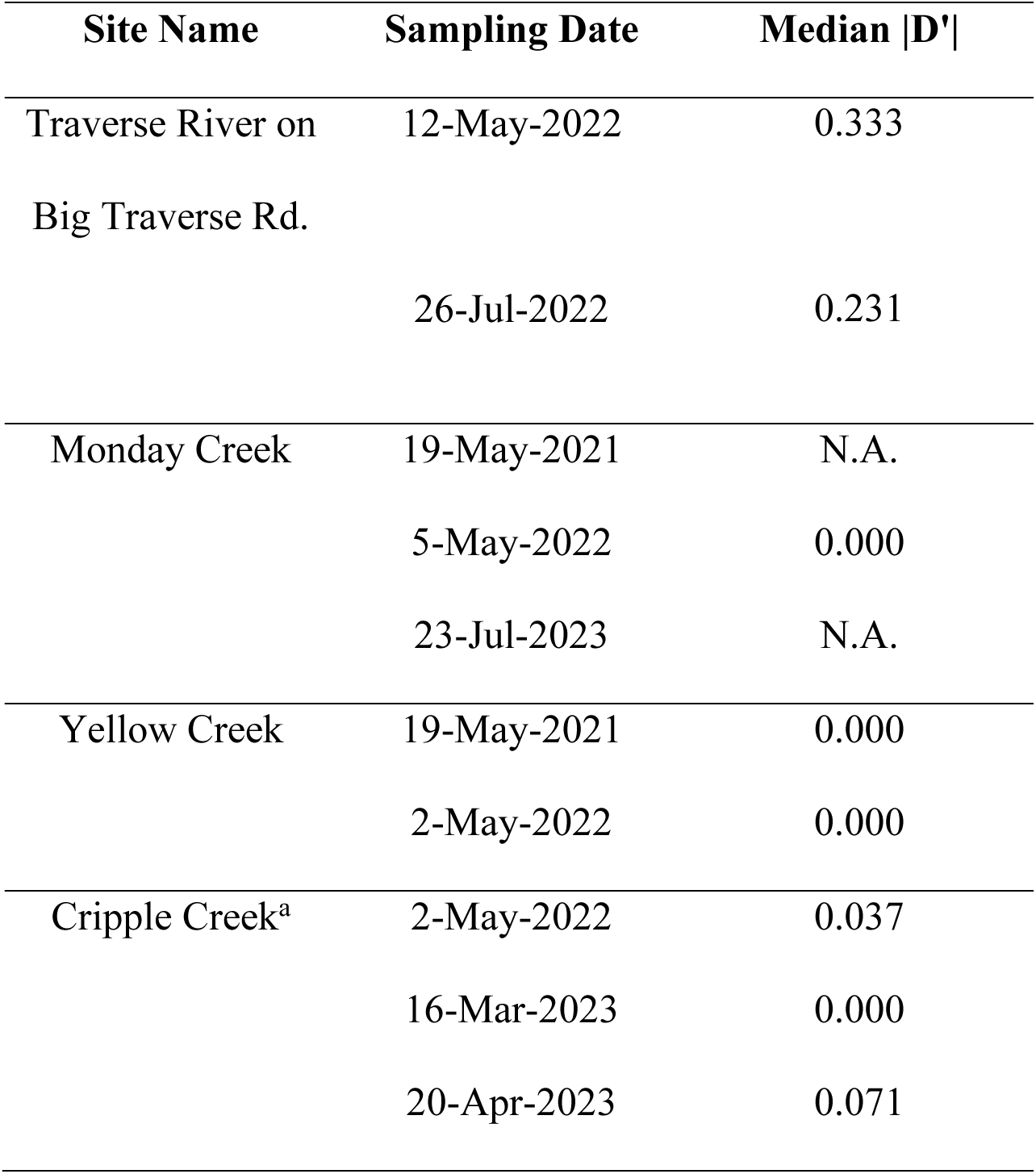
Median per-locus linkage disequilibrium (**|**D’|) for each time point. At some time points, all loci were fixed and **|**D’| could not be calculated. In these situations, the median **|**D’| is indicated as not applicable (“N.A.”).

**Table S6.**
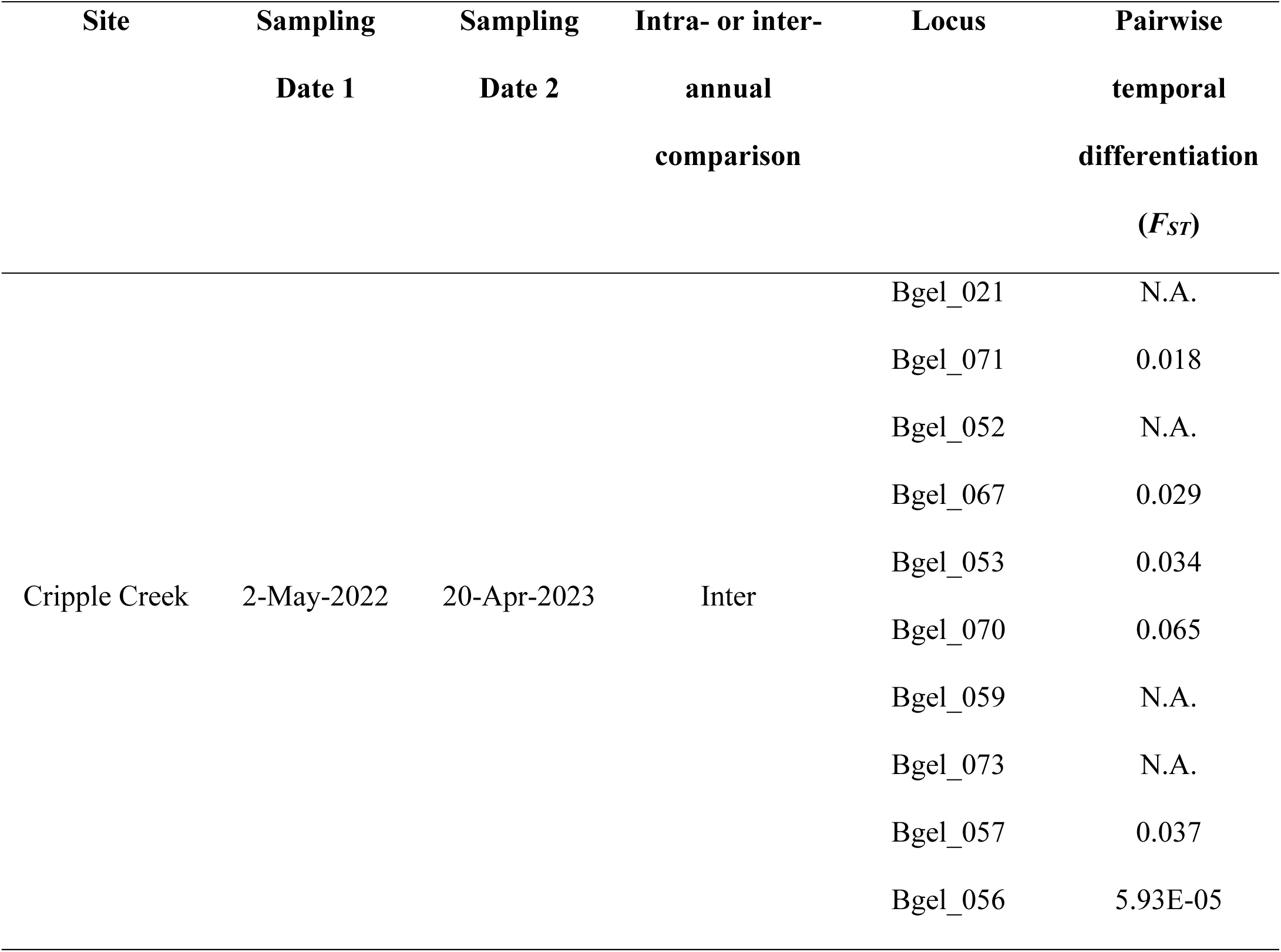
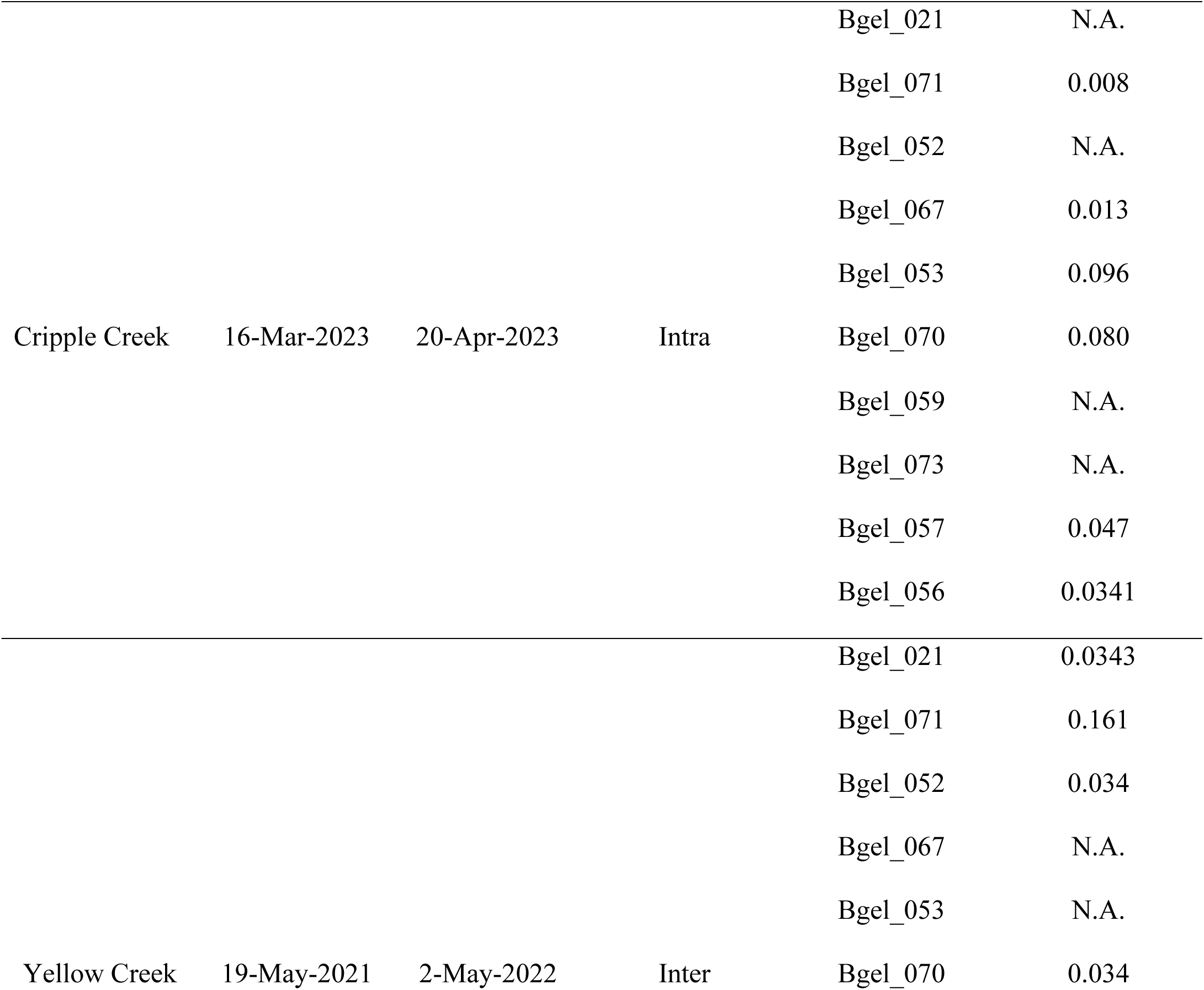
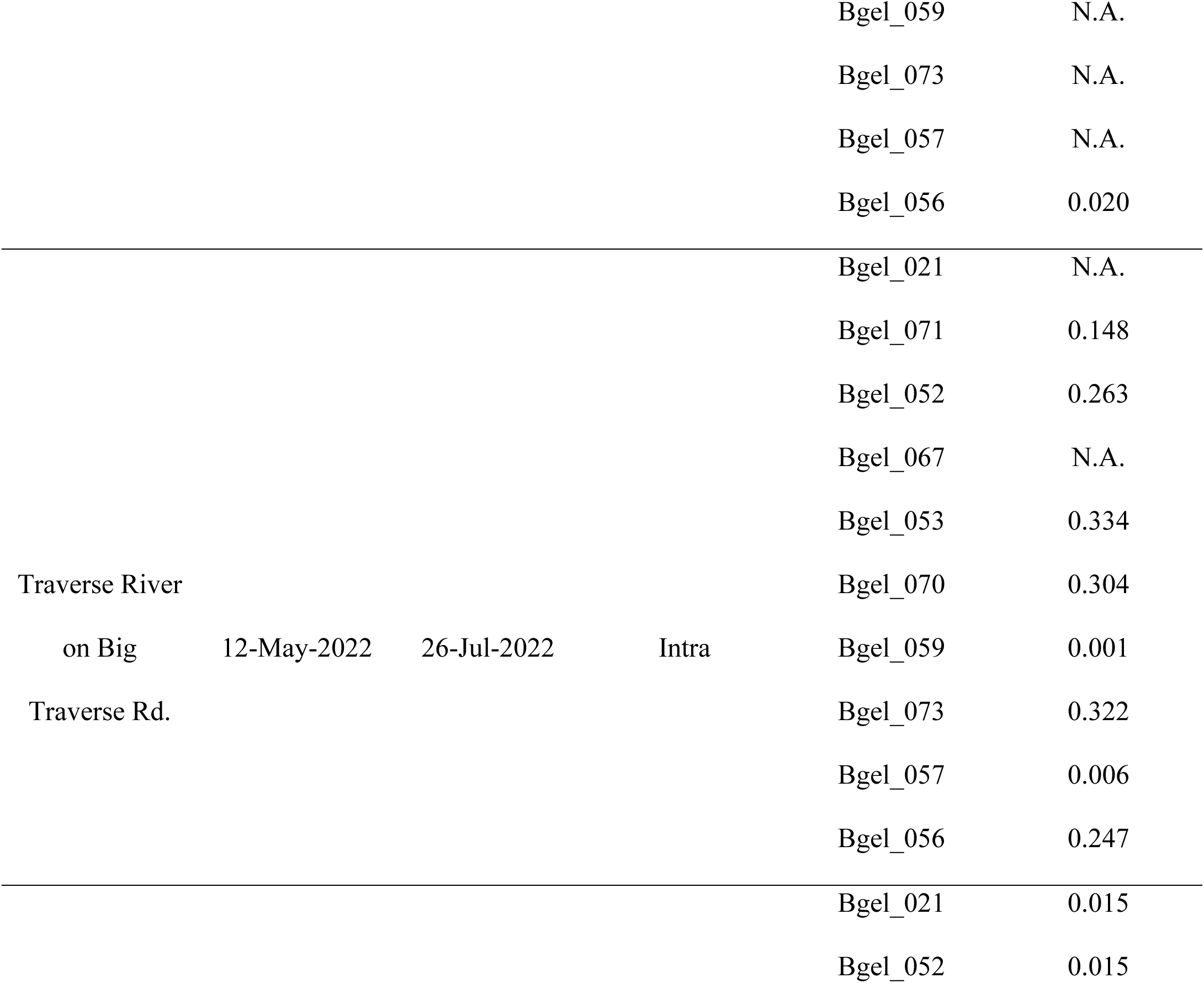
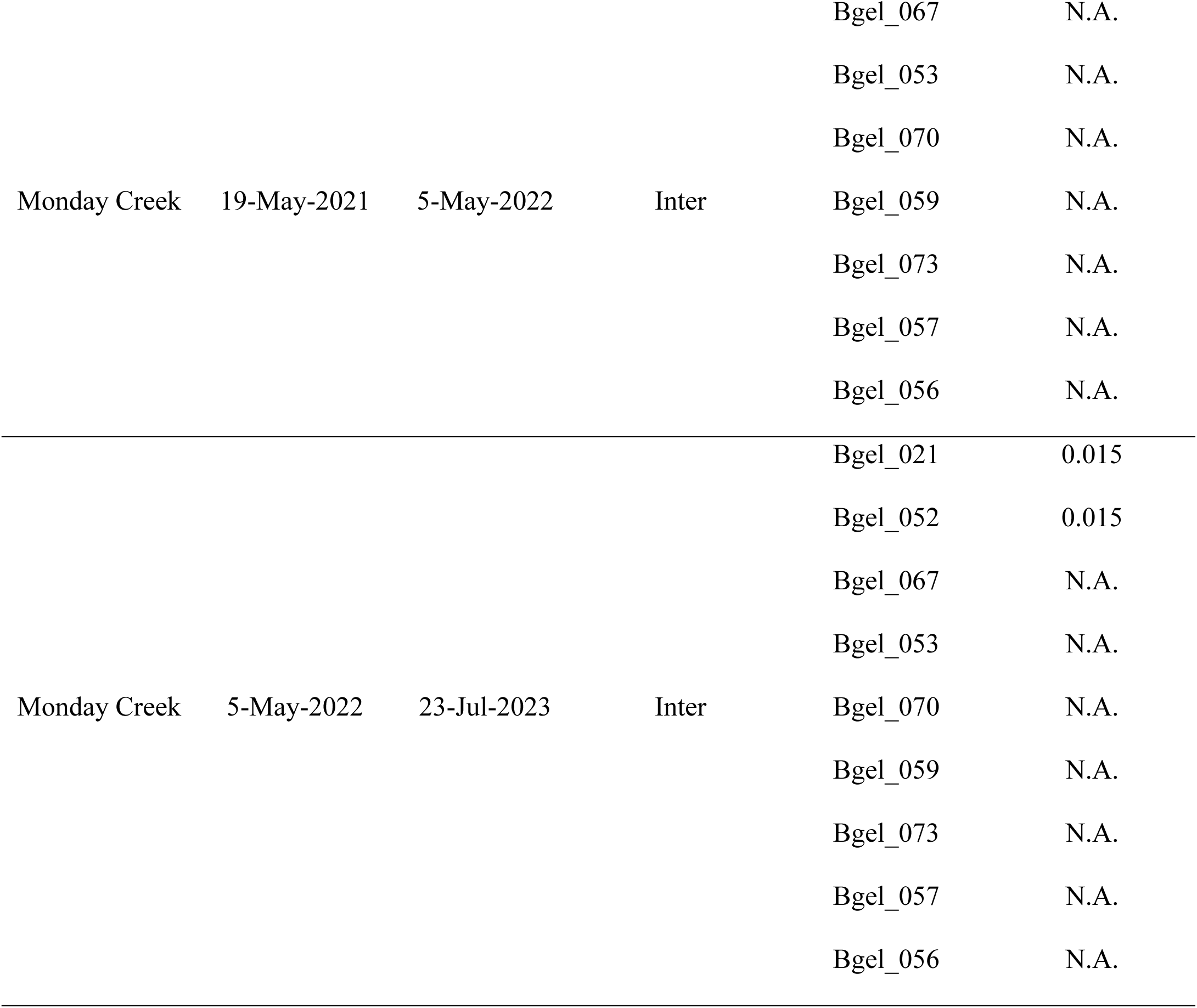
Pairwise temporal differentiation per locus at each site and time point (measured as *F_ST_*). Differentiation could not be measured if the locus was fixed at both time points. In these situations, the pairwise temporal differentiation is indicated as not applicable (“N.A.”).

**Table S7.** Freshwater red algal species observed as macroscopic gametophytic thalli at each time point sampled at AL-CRC.

| Date | Species present |
| --- | --- |
| May 2022 | *Audouinella* sp.<br>*Batrachospermum gelatinosum*<br>*Paralemanea* sp.<br>*Tuomeya americana* |
| March 2023 | *Batrachospermum gelatinosum*<br>*Paralemanea* sp.<br>*Tuomeya americana* |
| April 2023 | *Audouinella* sp.<br>*Batrachospermum gelatinosum*<br>*Paralemanea* sp.<br>*Kumanoa holtonii* |

**Figure S1.**
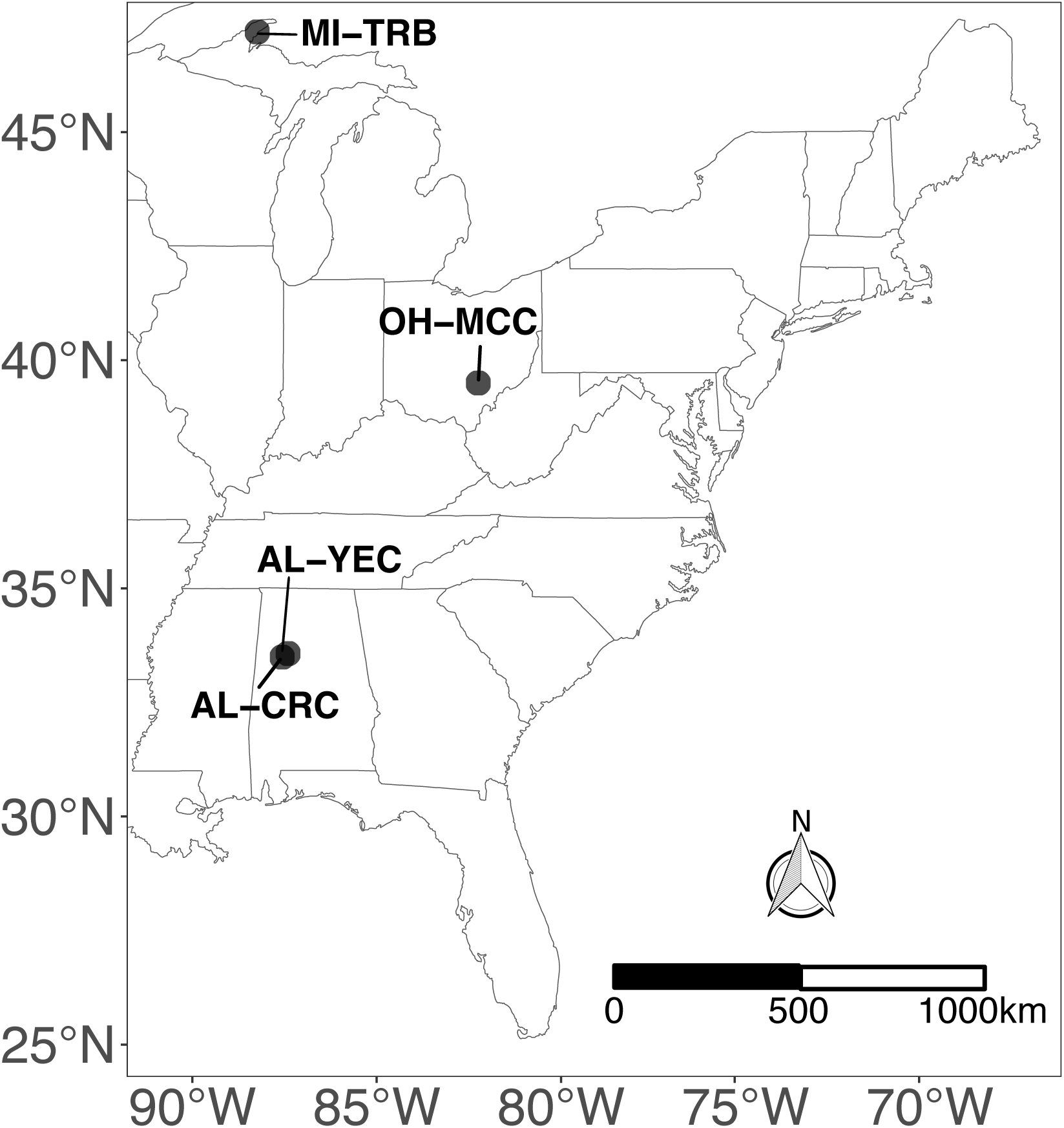
Map of four collection sites for *Batrachospermum gelatinosum.* GPS coordinates, physiochemical measurements, and sampling dates are in Table 2.

